# DKC1-mediated pseudouridylation of rRNA targets hnRNP A1 to sustain IRES-dependent translation and ATF4-driven metabolic adaptation

**DOI:** 10.1101/2024.12.12.628228

**Authors:** Anamika Gupta, Mohit Bansal, Jane Ding, Madhuparna Pandit, Sunil Sudarshan, Han-Fei Ding

## Abstract

The pseudouridine synthase DKC1 regulates internal ribosome entry site (IRES)-dependent translation and is upregulated in cancers by the MYC family of oncogenic transcription factors. We investigated the functional significance of DKC1 in MYCN-amplified neuroblastoma and its underlying mechanisms. A key function of DKC1 is to promote an ATF4-mediated gene expression program for amino acid metabolism and stress adaptation. We identified hnRNP A1, an IRES trans-acting factor, as a critical downstream mediator of DKC1 in sustaining ATF4 expression and IRES-dependent translation. We found that DKC1-mediated pseudouridylation at two specific 28S rRNA sites is essential for maintaining hnRNP A1 protein expression. Moreover, hnRNP A1 interacts with and stabilizes ATF4 mRNA, significantly increasing the protein expression of the ATF4 V1 variant, which contains an IRES element in its mRNA. Additionally, we found that cellular stress induces hnRNP A1, which is required for ATF4 induction under such conditions. Collectively, our study reveals a MYC-activated DKC1-hnRNP A1 axis that drives ATF4-mediated metabolic adaptation, supporting cancer cell survival under metabolic stress during cancer development.

## Introduction

The DKC1 gene encodes an evolutionarily conserved protein, dyskerin (hereafter referred to as DKC1) ^1^. Mutations in the DKC1 gene cause X-linked dyskeratosis congenita (X-DC), a genetic disorder characterized by defects in proliferating tissues, leading to symptoms such as bone marrow failure, immunodeficiency, abnormal skin pigmentation, nail dystrophy, stem cell defects, premature aging, and an increased susceptibility to cancer ^1–3^. Hypomorphic Dkc1 mutant mice, which express Dkc1 at approximately 30% of normal levels, recapitulate key symptoms of X-DC, including bone marrow failure and increased cancer susceptibility ^4^.

DKC1 is a pseudouridine synthase that catalyzes the conversion of uridine to pseudouridine through base rotation ^5–8^. Unlike other members of the pseudouridine synthase family, DKC1 functions within a complex of ribonucleoproteins and box H/ACA small nucleolar RNAs (snoRNAs). Each snoRNA within this complex acts as a guide, using base complementarity to select the target RNA and the specific uridine for pseudouridylation ^7,9^. The primary targets of DKC1-mediated pseudouridylation are ribosome RNAs (rRNAs) and spliceosomal small nuclear RNAs (snRNAs). Consequently, DKC1 plays a critical role in the biogenesis of ribosomes and spliceosomes ^10^.

A 2006 study provided the first evidence of a role for DKC1 in regulating internal ribosome entry site (IRES)-dependent translation. Cells from X-DC patients and Dkc1 hypomorphic mutant mice exhibited reduced translation of mRNAs containing IRES elements, such as those encoding p27Kip1, BCL-XL, and XIAP ^11^. Subsequent studies demonstrated that reduced DKC1 levels also impaired IRES-dependent translation of p53 mRNA ^12,13^. Moreover, it was shown that a reduction in DKC1-mediated rRNA pseudouridylation decreased ribosome affinity to a viral IRES element ^14^, suggesting that DKC1-mediated pseudouridylation of rRNA promotes IRES-dependent translation. However, the detailed mechanisms underlying this process remain poorly understood.

We investigated the role of DKC1 in cancer, motivated by genetic evidence linking DKC1 mutations to proliferative defects ^1–3,15^. In addition, DKC1 has been identified as a direct target gene of the MYC family of oncogenic transcription factors, including MYC ^16^ and MYCN ^17^, which drive the development of many cancers, such as neuroblastoma, by promoting cell proliferation and metabolic reprogramming ^18–21^. Neuroblastoma, a pediatric cancer of the sympathetic nervous system, accounts for 15% of childhood cancer-related deaths ^22,23^. DKC1 is highly expressed in neuroblastoma and is essential for the tumorigenic growth of neuroblastoma cell lines^17^. Our investigation revealed that a key function of DKC1 is to sustain both steady-state and stress-induced expression of ATF4, the master transcriptional regulator of amino acid metabolism and the integrated stress response (ISR) ^24–27^. This function of DKC1 is mediated through pseudouridylation of 28S rRNA, which promotes hnRNP A1 protein synthesis to drive IRES-dependent translation and ATF4 expression. Collectively, our study uncovers a MYC-activated DKC1 pseudouridylation program that facilitates cancer metabolic reprogramming and stress adaptation through hnRNP A1-mediated IRES-dependent translation.

## Results

### DKC1 sustains the expression of ATF4 and key genes involved in amino acid metabolism

We analyzed RNA-seq datasets from independent neuroblastoma patient cohorts ^28,29^, revealing that high DKC1 mRNA levels are significantly associated with advanced neuroblastoma stages and poor patient prognosis (Figures S1A and S1B). Consistent with previous findings ^17^, knockdown of DKC1 expression by shRNA (shDKC1) in neuroblastoma cell lines inhibited cell proliferation and xenograft growth, prolonging the survival of tumor-bearing mice (Figures S1C-1F). Conversely, DKC1 overexpression in neuroblastoma cell lines significantly accelerated their growth (Figures S1G and S1H). These results provide a strong rationale for using neuroblastoma as a model system to explore DKC1-mediated RNA pseudouridylation in cancer and its underlying mechanisms.

To gain molecular insight into the growth-promoting function of DKC1 in neuroblastoma, we performed RNA-seq analysis on MYCN-amplified neuroblastoma BE(2)-C cells with shRNA-mediated DKC1 knockdown (Supplementary Table 1). A total of 531 genes (≥ −1.50 fold, *P* < 0.05) were downregulated following DKC1 knockdown (Supplementary Table 1). Gene ontology (GO) analysis revealed that the downregulated genes were significantly enriched for GO terms related to amino acid and pyrimidine nucleotide metabolic processes (Figure 1A and Supplementary Table 2). These downregulated genes include those encoding ATF4, amino acid synthesis enzymes (e.g., ASNS, PSAT1, SHMT2), and amino acid transporters (e.g., SLC7A5 and SLC7A11) (Figure 1B and Supplementary Table 2). Gene set enrichment analysis (GSEA) further revealed that DKC1 knockdown reduced mRNA expression of genes involved in the cellular response to amino acid starvation (AAR) mediated by the eIF2α kinase EIF2AK4 (GCN2) ^24–27^ (Figure 1C). To validate the shDKC1 RNA-seq data, we performed RNA-seq analysis on BE(2)-C cells treated with a DKC1 inhibitor, pyrazofurin ^30^ and obtained similar results: DKC1 inhibition downregulated ATF4 and genes involved in amino acid synthesis and the GCN2-mediated AAR (Figures S2A and S2B and Supplementary Table 3). We further confirmed the RNA-seq findings by analyzing published proteomics data ^31^, showing that DKC1 knockdown decreased the expression of proteins involved in amino acid synthesis and transport, and one-carbon metabolism, including ASNS, MTHFD2, PHGDH, PSAT1, SHMT2, and SLC7A5 (Figure S2C and Supplementary Table 4).

**Figure 1:**
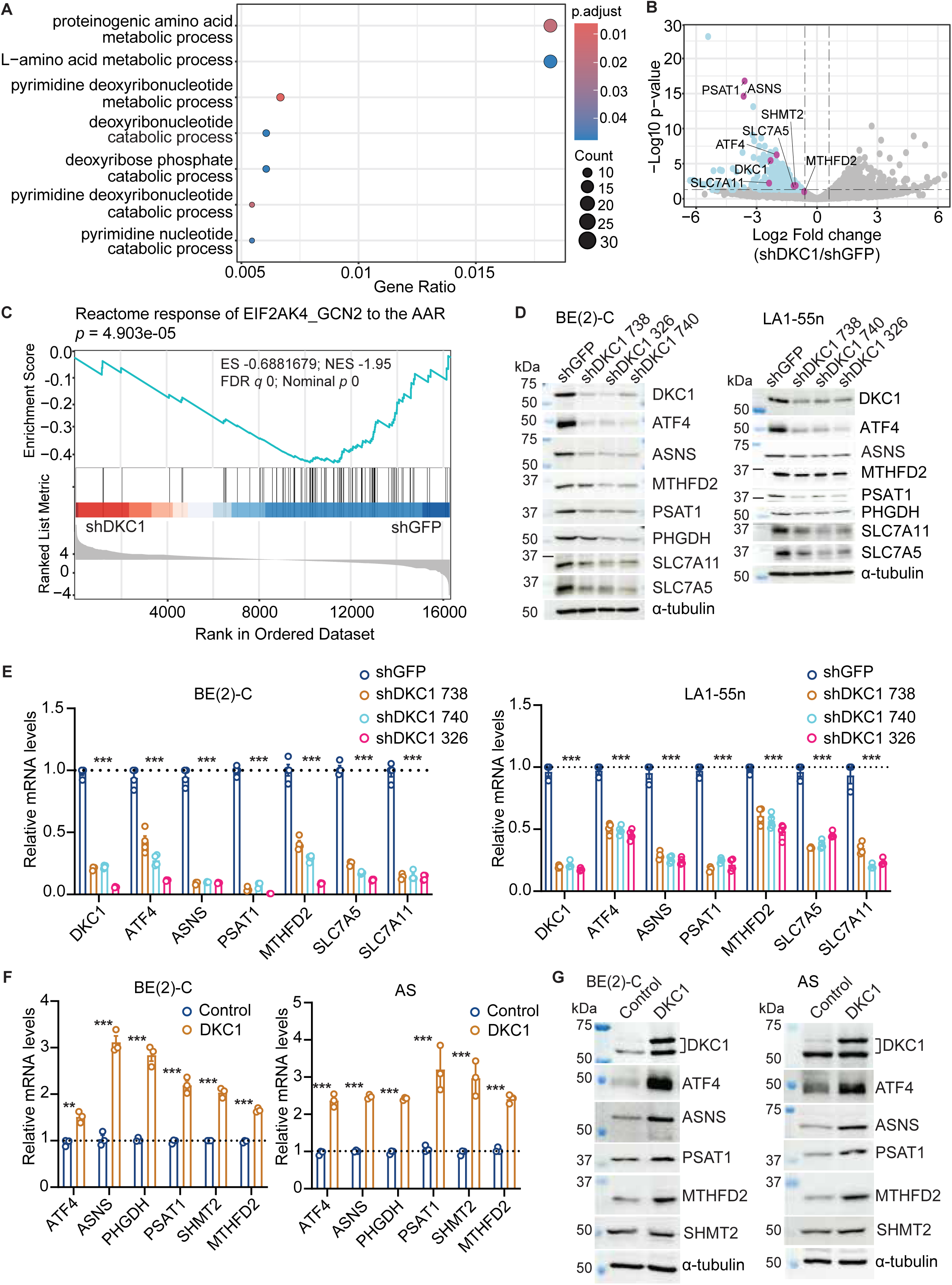
DKC1 sustains the expression of ATF4 and genes involved in amino acid metabolism. **(A)** GO analysis of RNA-seq data showing top biological processes for genes downregulated (≥ −1.5 fold) in neuroblastoma BE(2)-C cells following DKC1 knockdown. **(B)** Volcano plot showing downregulation of genes involved in amino acid metabolism following DKC1 knockdown. **(C)** GSEA of RNA-seq data showing downregulation of genes involved in the EIF2AK4_GCN2 response to amino acid deficiency following DKC1 knockdown. **(D-E)** Immunoblotting (**D**) and qRT-PCR (**E**) showing downregulation of genes involved in amino acid metabolism following DKC1 knockdown in neuroblastoma BE(2)-C and LA1-55n cells. **(F-G)** qRT-PCR (**F**) and immunoblotting **(G)** showing DKC1 overexpression upregulated genes involved in amino acid metabolism in neuroblastoma BE(2)-C and SK-N-AS cells. qRT-PCR data (**E and F**) are presented as mean ± SEM (n = 4) and analyzed using two-way ANOVA, \*\*\**P* <0.001.

We performed quantitative RT-PCR (qRT-PCR) and immunoblot analyses to validate the RNA-seq and proteomics findings. DKC1 knockdown using multiple shDKC1 constructs (Figures 1D and 1E) or inhibition (Figures S2D–S2G) led to reduced mRNA and protein expression of the same set of genes. Conversely, DKC1 overexpression resulted in increased mRNA and protein expression of these genes (Figures 1F and 1G).

To determine whether DKC1 regulation of ATF4 is physiologically relevant, we assessed the impact of DKC1 inhibition on cellular stress responses using models of endoplasmic reticulum (ER) stress and the AAR. We treated BE(2)-C and HeLa cells with tunicamycin (an ER stress inducer) or histidinol (HisOH, an AAR inducer) ^32,33^ in the absence or presence of the DKC1 inhibitor pyrazofurin. In control cells, tunicamycin or HisOH treatment induced ATF4, and this induction was significantly reduced in cells treated with pyrazofurin (Figure S2H).

Collectively, these data demonstrate that DKC1 plays a crucial role in maintaining the expression of ATF4 and genes essential for amino acid biosynthesis and transport, providing a molecular mechanism for the functional importance of DKC1 in neuroblastoma.

### DKC1 regulates amino acid metabolism

In light of the gene expression data, we performed targeted metabolomics to investigate the effect of DKC1 inhibition on cellular metabolism. Metabolomics profiling of BE(2)-C cells treated with the DKC1 inhibitor pyrazofurin revealed significant changes (p < 0.05) in the levels of 131 metabolites (Supplementary Table 5). Pathway analysis of the downregulated metabolites using MetaboAnalyst ^34^ showed significant enrichment of metabolic pathways related to amino acid metabolism, including those for alanine, glutamate, methionine, glycine and serine, and aspartate (Figure 2A). An examination of individual amino acids indicated broad changes in their levels, consistent with the gene expression data and highlighting a key role of DKC1 in regulating amino acid metabolism. For instance, the levels of essential amino acids such as phenylalanine, threonine, tryptophan, and valine were significantly reduced following DKC1 inhibition (Figure 2B), potentially due to the downregulation of SLC7A5 (Figures 1B, 1D, and 1E), a transporter responsible for their import ^35,36^. We further analyzed the serine-glycine synthesis pathway, which uses the glycolytic intermediate 3-phosphoglycerate (3PG) to generate serine and glycine (Figure 2C). DKC1 inhibition reduced the expression of pathway enzymes PHGDH, PSAT1, and SHMT2 (Figures S2D-S2G), resulting in a significant decrease in glycine levels and an accumulation of upstream intermediates, including serine and 3PG (Figure 2D). Collectively, the metabolomics and gene expression data underscore the critical role of DKC1 in regulating amino acid metabolism in neuroblastoma cells.

**Figure 2.**
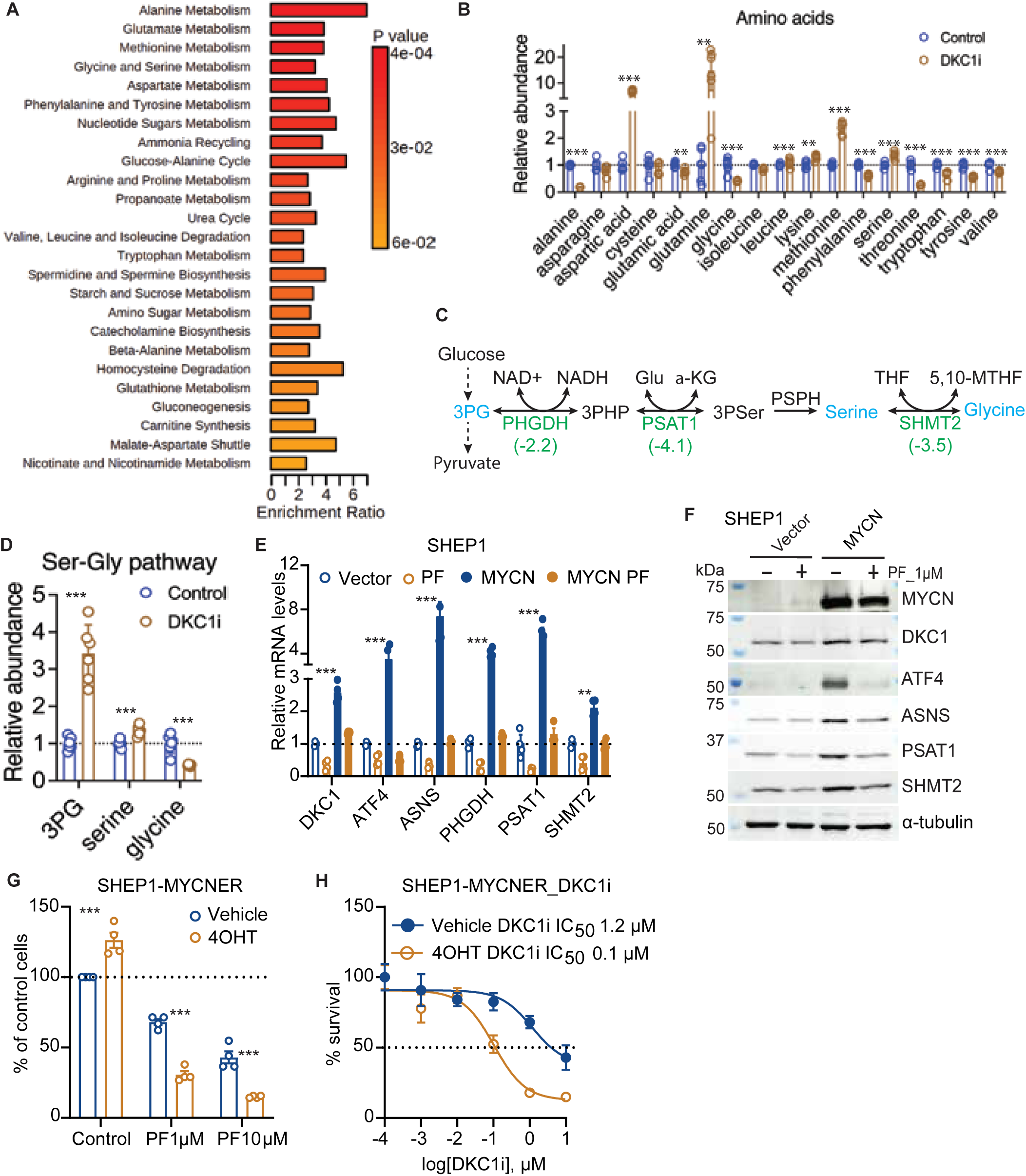
DKC1 is a downstream mediator of MYCN in regulation of amino acid metabolism. **(A)** Enriched metabolic pathways identified through metabolite set enrichment analysis (MSEA) of downregulated metabolites in BE(2)-C cells treated with the DKC1 inhibitor pyrazofurin 1µM for 72 h. **(B)** Relative levels of amino acids in BE(2)-C cells following DKC1 inhibition. **(C)** Schematic of the serine-glycine synthesis pathway with indicated enzymes and metabolites. Fold changes (numbers in parentheses) in mRNA expression of relevant enzymes were from qRT-PCR. (**D**) Relative levels of the serine-glycine pathway metabolites. (**B, D**) Data represent mean ± SEM (n = 6) and were analyzed by unpaired, two-tailed Student’s t-test. **(E-F)** qRT-PCR (**E**) and Immunoblotting (**F**) showing DKC1 inhibition abrogated MYCN upregulation of genes for amino acid synthesis in SHEP1 cells. (**G**) MYCN activation by 4OHT promoted cell proliferation (vehicle) but sensitized cells to DKC1 inhibition (PF). (**H**) IC_50_ analysis of the DKC1 inhibitor pyrazofurin (DKC1i) with or without MYCN activation by 4OHT for 24 h, followed by treatment with different concentrations of DKC1i for 24h. Data (**E, G, and H**) are presented as mean ± SEM (n = 4) and were analyzed using two-way ANOVA. \*\**P* < 0.01, \*\*\**P* <0.001.

### DKC1 is a downstream mediator of MYCN in sustaining the expression of ATF4 and genes involved in amino acid metabolism

DKC1 has been identified as a direct target gene of MYCN in neuroblastoma ^17^. Consistent with this finding, RNA-seq data from the SEQC neuroblastoma patient cohort ^28^ demonstrated a strong positive correlation in mRNA expression between DKC1 and MYCN in neuroblastoma tumors (Figure S3A). We confirmed the ability of MYCN to upregulate DKC1 expression through qRT-PCR and immunoblot analyses (Figures 2E, 2F, S3B, and S3C).

Previously, we showed that a key function of MYCN in neuroblastoma metabolic reprogramming is to transcriptionally upregulate amino acid synthesis enzymes via ATF4 ^37^. This function of MYCN was abrogated by DKC1 inhibition (Figures 2E and 2F) or knockdown (Figures S3B and S3C). Furthermore, DKC1 inhibition impaired the ability of MYCN to promote neuroblastoma cell proliferation (Figures 2G and S3D), and MYCN overexpression sensitized neuroblastoma cells to the DKC1 inhibitor pyrazofurin, resulting in a 12-fold decrease in its IC_50_ (Figure 2H). Altogether, these findings suggest that DKC1 is a critical downstream mediator of MYCN, driving ATF4 expression and amino acid synthesis to sustain neuroblastoma cell proliferation. Our results underscore the functional significance of DKC1 upregulation by MYCN in neuroblastoma.

### DKC1 targets the IRES trans-acting factor hnRNP A1

To further investigate into the mechanism of DKC1 action, we aimed to identify its downstream effector(s) involved in the regulation of IRES-dependent translation. We used Venn Diagram analysis to integrate three gene lists: (1) genes downregulated by DKC1 inhibition (≥ −1.50-Log2 fold, adjusted *P* < 0.01), (2) genes positively correlated with DKC1 expression in neuroblastoma tumors (R > 0.7, *P* < 1.14E-72) ^28^, and (3) genes encoding IRES trans-acting factors (ITAFs) ^38^ (http://www.iresite.org/IRESite_web.php?page=ITAFs_of_cellular_IRESs). Our analysis pinpointed a single candidate that satisfied all criteria: hnRNP A1, encoded by the HNRNPA1 gene. Multiple lines of correlative evidence support the notion that hnRNP A1 is a potential DKC1 effector. First, both DKC1 ^11,39,40^ and hnRNP A1 ^41,42^ are known regulators of IRES-mediated translation. Second, HNRNPA1 mRNA levels positively correlate with DKC1 mRNA levels in neuroblastoma tumors (Figure 3B). Third, DKC1 and hnRNP A1 share a similar phenotype in neuroblastoma: 1) Higher HNRNPA1 mRNA expression is significantly associated with higher MYCN mRNA expression, advanced neuroblastoma stages, and poor patient prognosis (Figures S4A-S4C); 2) Overexpression of hnRNP A1 stimulated neuroblastoma cell growth (Figures 3C and 3D); and 3) knockdown of hnRNP A1 expression using various shHNRNPA1 constructs (Figure 3E) inhibited neuroblastoma cell growth in culture (Figure 3F) and in immunodeficient mice (Figure 3G), leading prolonged survival of xenograft-bearding mice (Figure 3H). Thus, like DKC1, high hnRNP A1 expression is required for the proliferation and tumorigenicity of neuroblastoma cells.

**Figure 3.**
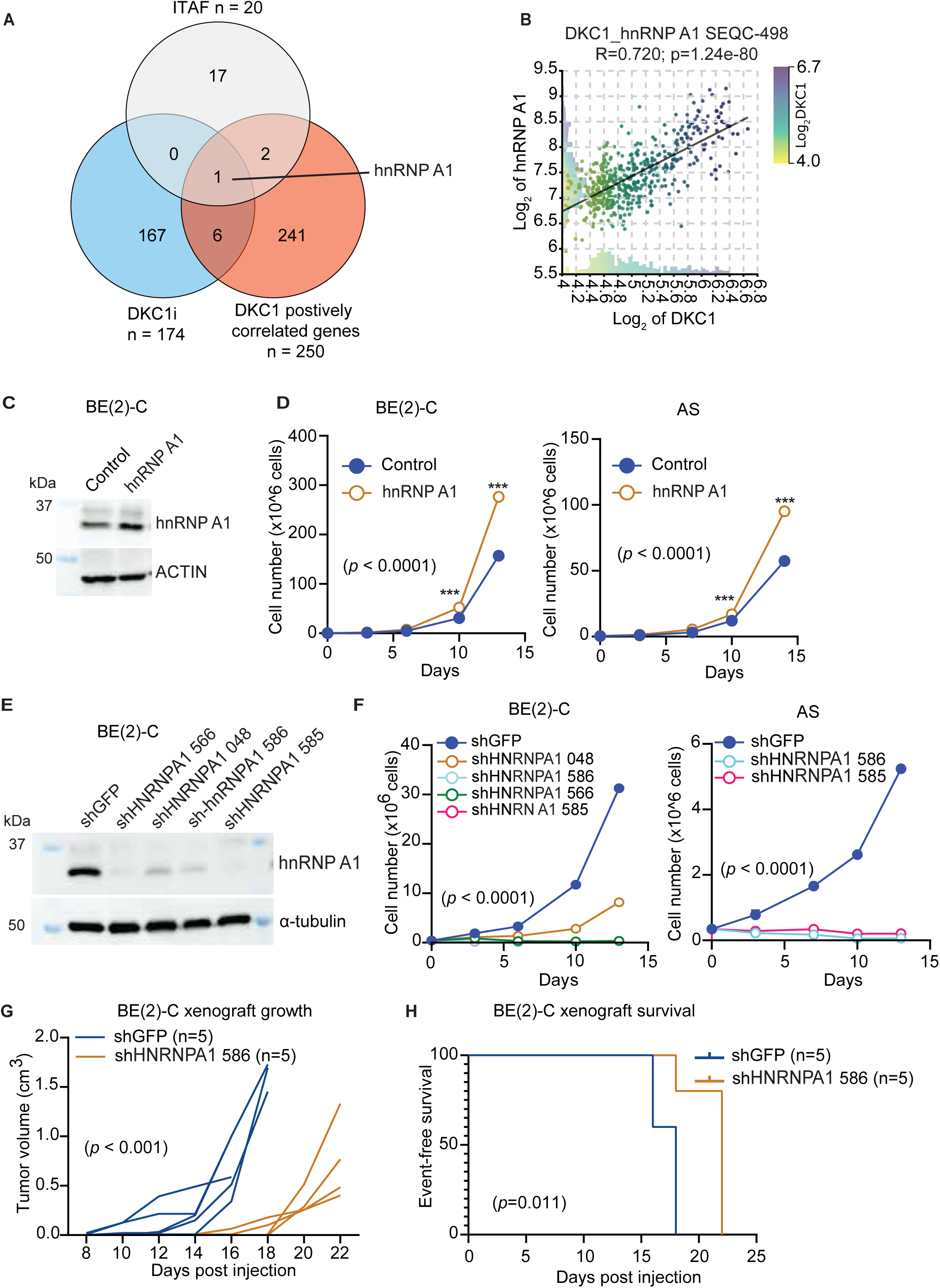
DKC1 targets the IRES trans-acting factor hnRNP A1. **(A)** Venn diagram illustration of hnRNP A1 as a shared gene among the lists of DKC1 positively correlated genes in the SEQC neuroblastoma patient dataset, genes downregulated by DKC1 inhibition (DKC1i), and IRES trans-acting factors. **(B)** Positive correlation in mRNA expression between DKC1 and HNRNPA1 in primary neuroblastoma tumors (the SEQC cohort, n = 498). R (correlation) and p values are indicated. **(C)** Immunoblotting of hnRNP A1 in neuroblastoma BE(2)-C cells with hnRNP A1 overexpression. **(D)** hnRNP A1 overexpression promoted the proliferation of BE(2)-C and SK-N-AS neuroblastoma cells. **(E)** Immunoblotting of hnRNP A1 in neuroblastoma BE(2)-C cells with hnRNP A1 knockdown. **(F)** hnRNP A1 knockdown reduced the proliferation of BE(2)-C and SK-N-AS neuroblastoma cells. **(G-H)** hnRNP A1 knockdown impeded BE(2)-C xenograft growth **(G)** and prolonged the event-free survival of xenograft-bearing mice. Cell growth data **(D** and **F)** are presented as mean ± SEM (n = 4) and quantitative data **(D, F,** and **G)** were analyzed using two-way ANOVA. \*\*\**P* <0.001.

We obtained directing evidence indicating that hnRNP A1 is a downstream target DKC1. Knockdown of DKC1 (Figures 4A and 4B) or its inhibition (Figures 4C and 4D) significantly reduced both HNRNPA1 mRNA and hnRNP A1 protein expression in neuroblastoma cell lines. Similar results were observed with DKC1 knockdown in HeLa cells (Figures 4A and 4B), demonstrating that the regulation of hnRNP A1 expression by DKC1 is not cell-type specific.

**Figure 4.**
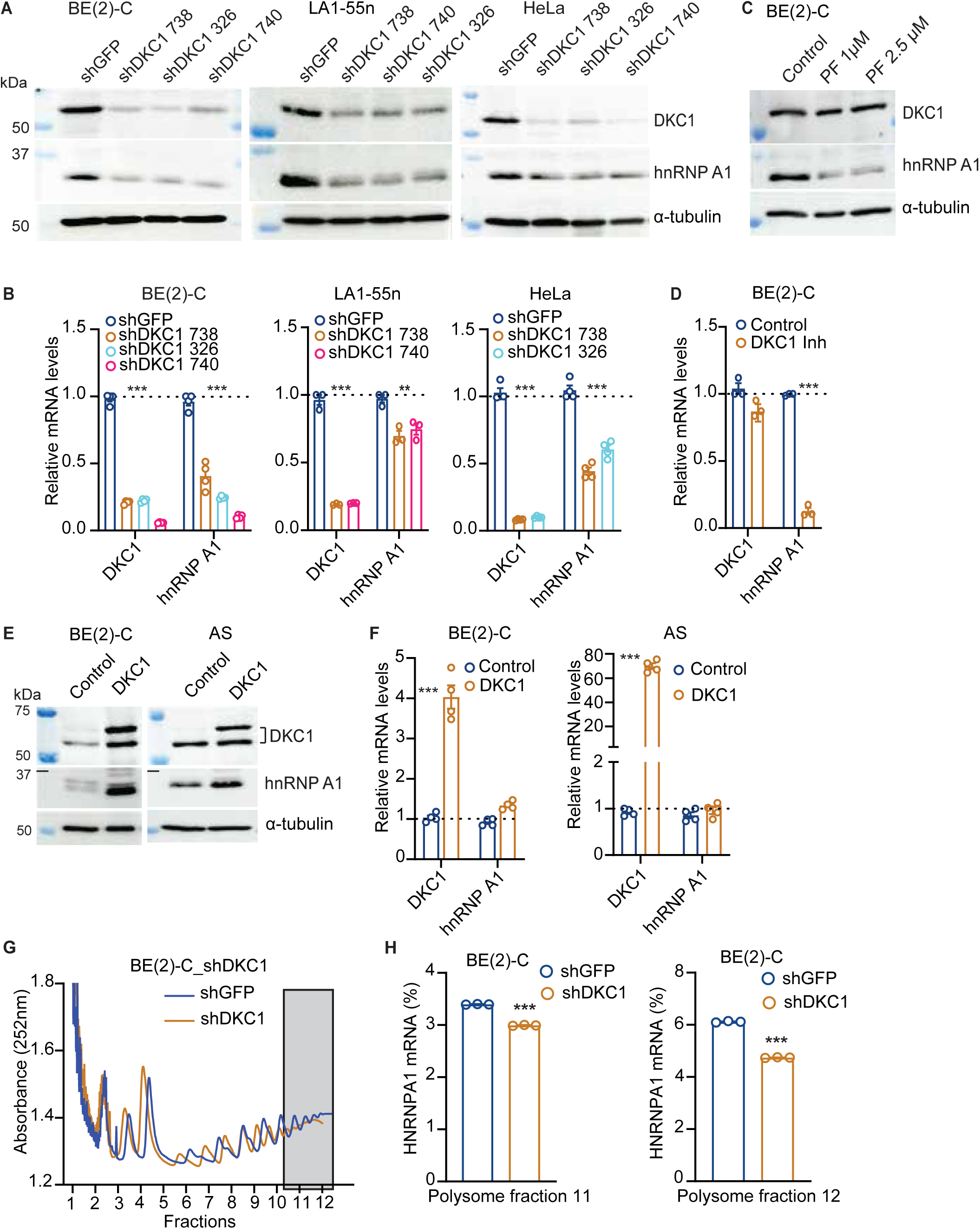
DKC1 promotes HNRNPA1 mRNA translation. (**A-B)** Immunoblotting (**A**) and qRT-PCR **(B)** showing the downregulation of hnRNP A1 protein and mRNA expression following DKC1 knockdown in the indicated cell lines. (**C-D)** Immunoblotting (**C**) and qRT-PCR **(D)** showing the downregulation of hnRNP A1 protein and mRNA expression following DKC1 inhibition in BE(2)-C cells. **(E-F)** Immunoblotting (**E**) and qRT-PCR **(F)** showing the upregulation of hnRNP A1 protein expression but not HNRNPA1 mRNA expression following DKC1 overexpression in BE(2)-C and SK-N-AS cells. **(G-H)** Polysome profiling (**G**) and qRT-PCR analysis of polysome fractions (**H**) showing a significant reduction in the levels of polysome-associated HNRNPA1 mRNA. Data represent two independent experiments and are presented as the mean ± SEM of 3 technical replicates. qRT-PCR data were analyzed using two-way ANOVA. \*\**P* < 0.01, \*\*\**P* < 0.001.

Notably, DKC1 overexpression increased hnRNP A1 protein levels without significantly affecting its mRNA expression (Figures 4E and 4F), suggesting that DKC1 regulates HNRNPA1 mRNA translation. To investigate this possibility, we performed polysome profiling, as actively translating mRNAs are associated with higher numbers of ribosomes (polysomes) ^43^. Knockdown of DKC1 triggered a shift in the distribution of HNRNPA1 mRNAs to lower numbers of polysomes, significantly reducing the levels of HNRNPA1 mRNAs associated with the highest numbers of polysomes (Figure 4F, grey boxed area). This was confirmed by qRT-PCR analysis of polysome fractions (Figure 4H). DKC1 knockdown had no effect on the levels of polysome-associated GAPDH mRNA (Figure S4D). These findings suggest that DKC1 upregulates hnRNP A1 expression by promoting its mRNA translation.

### hnRNPA 1 is a downstream mediator of DKC1 in sustaining ATF4 expression

As shown above, a key function of DKC1 is to sustain ATF4 expression. Therefore, we investigated the functional relationship between DKC1 and hnRNP A1 in regulating ATF4 expression. Overexpression of hnRNP A1 completely reversed the negative effect of DKC1 knockdown on ATF4 expression (Figure 5A) and cell growth (Figure 5B) and conferred resistance to DKC1 inhibition (Figure S5A). These findings suggest that hnRNP A1 acts downstream of DKC1 in sustaining ATF4 expression and neuroblastoma cell proliferation. Interestingly, when hnRNP A1 was overexpressed, shDKC1 constructs reduced DKC1 mRNA expression (Figure S5B) but failed to decrease DKC1 protein levels (Figure 5A), suggesting that hnRNP A1 overexpression stabilizes the DKC1 protein. Indeed, we found that both DKC1 and hnRNP A1 are nuclear proteins (Figure S5C) and could be co-immunoprecipitated with antibodies against DKC1 or hnRNP A1 (Figures S5D and S5E). These observations suggest that DKC1 and hnRNP A1 can form a complex, providing a mechanism by which hnRNP A1 stabilizes DKC1KDC1 protein.

**Figure 5.**
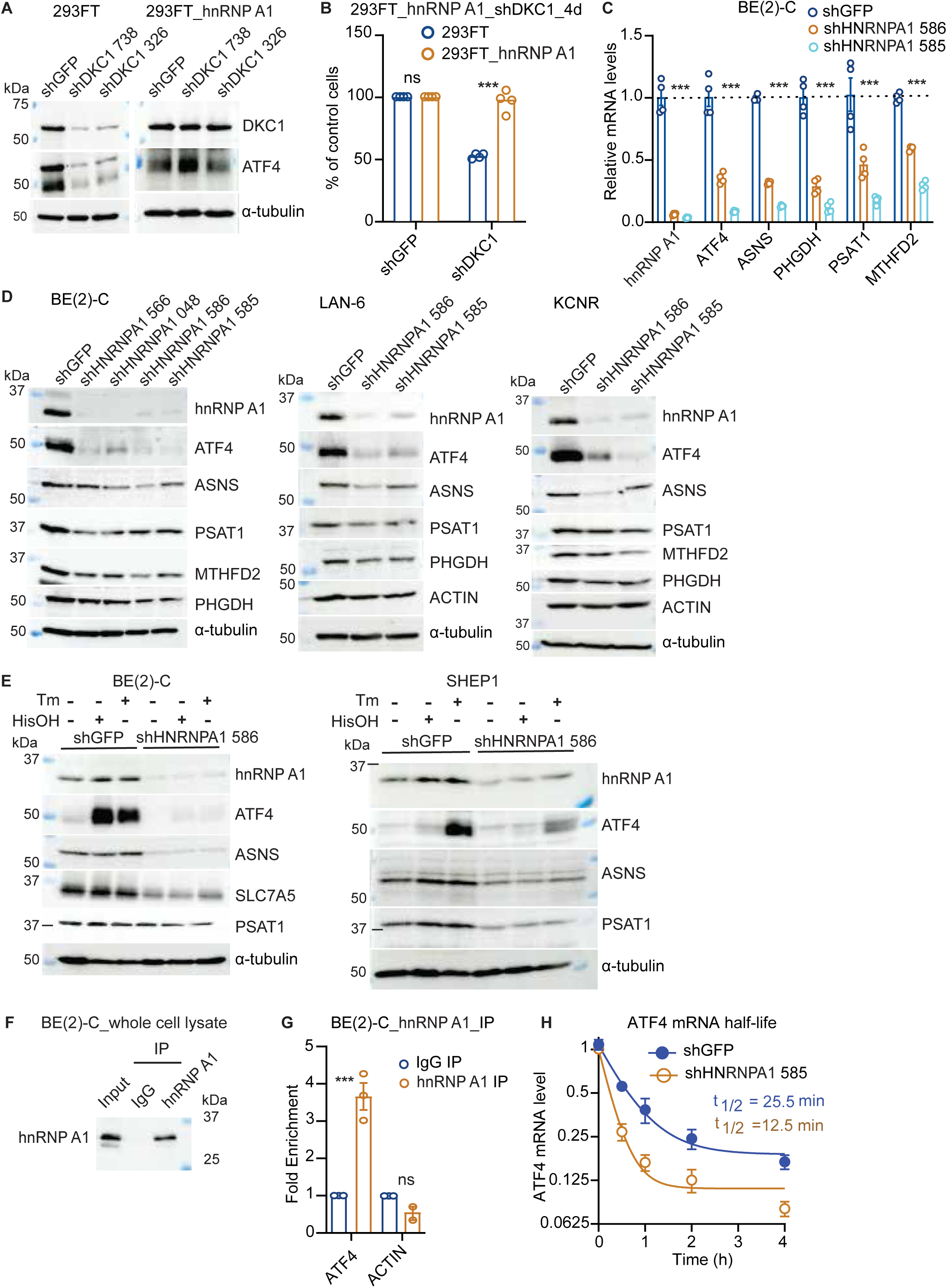
hnRNPA 1 sustains ATF4 expression and stress responses. **(A)** hnRNP A1 overexpression in 293FT cells alleviated the inhibitory effect of DKC1 knockdown on ATF4 protein expression. **(B)** hnRNP A1 overexpression alleviated the inhibitory effect of DKC1 knockdown on cell proliferation. Cell growth data are presented as mean ± SEM (n = 4) and analyzed using two-way ANOVA. **(C-D)** qRT-PCR (**C**) and immunoblotting (**D**) showing the downregulation of mRNA and protein expression of genes involved in amino acid metabolism following hnRNP A1 knockdown in the indicated neuroblastoma cell lines. qRT-PCR data are presented as mean ± SEM (n = 4) and analyzed using 2-way ANOVA. **(E)** Immunoblotting showing hnRNP A1 knockdown abrogated stress-induced upregulation of ATF4 protein and its downstream targets involved in amino acid metabolism in BE(2)-C and SHEP1 cells. HisOH, AAR inducer; Tm, ER stress inducer. **(F)** Immunoblotting of hnRNP A1 in control (IgG) and anti-hnRNP A1 immunoprecipitates. **(G)** qRT-PCR analysis of ATF4 and ACTIN mRNAs in the IgG and anti-hnRNP A1 immunoprecipitates. Data represent mean ± SEM (n = 2) and analyzed using 2-way ANOVA. **(H)** hnRNP A1 knockdown decreased ATF4 mRNA stability in BE(2)-C cells. RNA samples were collected at the indicated time points following addition of 5 µg/mL actinomycin D and analyzed by qRT-PCR. ATF4 mRNA levels were quantified against B2M and presented as the fraction of the initial level at time zero. Data represent mean ± SEM (n = 3). \*\*\**P* < 0.001; ns, not significant.

In further support of the model that hnRNP A1 acts downstream of DKC1, knockdown of hnRNP A1 expression using various shHNRNPA1 constructs fully recapitulated the effect of DKC1 knockdown, including downregulation of ATF4 and genes involved in amino acid metabolism at both mRNA and protein levels across multiple neuroblastoma cell lines (Figures 5C, 5D, and S5F). We also examined the effect of hnRNP A1 overexpression. The human HNRNPA1 gene produces two main mRNA variants, V1 (NM_002136.4) and V2 (NM_031157.4). Overexpression of both variants increased ATF4 protein expression (Figure S5G).

To determine the physiological relevance of hnRNP A1 regulation of ATF4, we evaluated the impact of hnRNP A1 knockdown on cellular stress responses. Control (shGFP) and hnRNP A1 knockdown BE(2)-C and SHEP1 cells were treated with tunicamycin (to induce ER stress) or HisOH (to activate the AAR). In control cells, treatment with tunicamycin or HisOH induced ATF4 and the expression of amino acid synthesis enzymes and transporters (Figure 5E). However, this induction was significantly reduced in hnRNP A1 knockdown cells (Figure 5E). Interestingly, both ER stress and the AAR also induced hnRNP A1 protein expression (Figure 5E), further supporting its critical role in cellular stress responses.

Next, we investigated the mechanisms by which hnRNP A1 sustains ATF4 expression. Given that hnRNP A1 is an RNA-binding protein (RBP) ^44,45^, we performed RNA immunoprecipitation using an antibody to hnRNP A1 (Figure 5F). Quantitative RT-PCR analysis showed a significant enrichment of ATF4 mRNA in the anti-hnRNP A1 precipitated RNA compared to the control (Figure 5G). To further validate these findings, we examined published RNA interactomes data generated through enhanced cross-linking immunoprecipitation (eCLIP) analysis of hundreds of human RBPs ^46^. The data confirmed the binding of hnRNP A1 to ATF4 mRNA in two different cancer cell lines, HepG2 and K562 (Figure S6). Moreover, knockdown of hnRNP A1 resulted in a two-fold reduction in the half-life of ATF4 mRNA (Figure 5H).

Taken together, these findings provide evidence that hnRNP A1 is a key downstream mediator of DKC1, promoting ATF4 expression by binding to and stabilizing its mRNA.

### hnRNP A1 promotes IRES-dependent ATF4 protein expression

Both DKC1 ^11,39^ and hnRNP A1 ^41,42,47–49^ are known to promote the translation of mRNAs containing IRES elements, including those encoding BCL-XL, CCND1, MYC, p27Kip1, and XIAP. As expected, DKC1 knockdown had a relatively small impact on the mRNA levels of BCL-XL, CCND1, and XIAP (Figure 6A), but markedly reduced their protein expression (Figure 6B and 6C). This effect of DKC1 knockdown was completely reversed by hnRNP A1 overexpression (Figure 6C). Similar results were observed with DKC1 inhibition (Figure 6D). Consistent with these findings, hnRNP A1 overexpression was sufficient to increase the protein expression of CCND1, BCL-XL, and XIAP (Figure 6E). These findings support the model that hnRNP A1 acts as a downstream mediator of DKC1 in regulating IRES-dependent translation.

**Figure 6.**
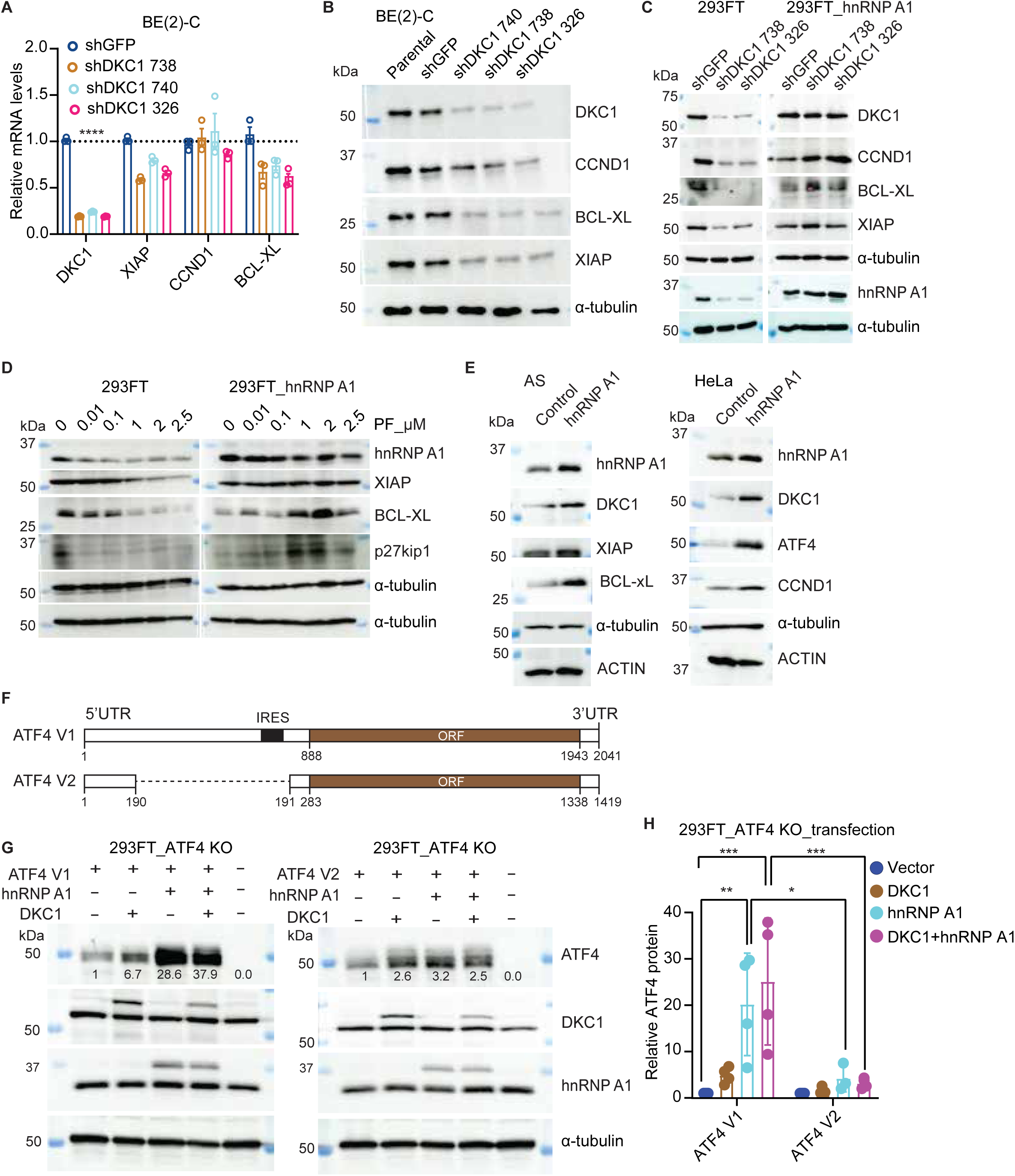
hnRNP A1 is a downstream mediator of DKC1 in regulation of IRES-dependent translation. **(A-B)** qRT-PCR (**A**) and immunoblotting **(B)** showing mRNA and protein expression of genes containing IRES elements following DKC1 knockdown. **(C-D)** hnRNP A1 overexpression alleviated the inhibitory effect of DKC1 knockdown (**C**) or inhibition (**D**) on IRES-dependent protein expression. **(E)** hnRNP A1 overexpression upregulated DKC1 and IRES-dependent protein expression. **(F)** Diagram of the ATF4 V1 and V2 mRNAs showing the main ORF and the IRES site in the 5’ UTR of ATF4 V1. **(G-H)** Immunoblotting (**G**) and quantification (**H**) of ATF4 variant protein expression 48 h after co-transfection of 293FT ATF4 KO cells with pGenLenti-ATF4 V1 or pCW57.1-ATF4 V2, in combination with plasmids expressing hnRNP A1, DKC1, or both hnRNP A1 and DKC1. ATF4 protein levels were quantified relative to α-tubulin. (**A, H**) data are presented as mean ± SEM (n = 4) and analyzed using two-way ANOVA. \**P* <0.05, \*\**P* < 0.01, \*\*\**P* <0.001.

The human ATF4 gene generates two main mRNA variants, V1 (NM_001675.4) and V2 (NM_182810.3), which encode the same protein but differ in their 5’ untranslated region (5’UTR). The V2 transcript lacks an internal segment present in the V1 transcript 5’UTR (Figure 6F). This internal segment in V1 reportedly contains an IRES element that mediates IRES-dependent translation of various reporter constructs ^50^ (Figure 6F). We hypothesized that DKC1 and hnRNP A1 may enhance IRES-dependent ATF4 mRNA translation and protein expression. To test this, 293FT ATF4 knockout cells were co-transfected with plasmids coding for either ATF4 V1 or V2, alongside DKC1- and/or HNRNPA1-expressing plasmids. Overexpression of DKC1 and hnRNP A1, alone or in combination, increased ATF4 V1 and V2 protein levels (Figures 6G and 6H). Importantly, the increase was significantly higher for ATF4 V1, particularly under hnRNP A1 overexpression. These results suggest that DKC1 and hnRNP A1 promote ATF4 protein expression predominantly through an IRES-dependent mechanism.

### DKC1-mediated 28S rRNA pseudouridylation targets hnRNP A1 to drive IRES-dependent translation and ATF4 expression

A previous study demonstrated that pseudouridylation at two specific sites in 28S rRNA, Ψ4331 and Ψ4966, is significantly reduced in ribosomes from patients with X-DC ^51^. To assess the impact of DKC1-dependent rRNA pseudouridylation on the regulation of hnRNP A1 and ATF4 expression, we analyzed rRNA samples collected on days 4 and 12 following DKC1 knockdown by Nanopore direct RNA sequencing. Consistent with previous findings, our analysis revealed a time-dependent reduction in pseudouridine levels at Ψ4331 and Ψ4966 in 28S rRNA (Figures 7A and 7B). In contrast, knockdown of another pseudouridine synthase, PUS7, had no effect on pseudouridylation at these sites (Figure S7A), confirming that pseudouridylation at these 28S rRNA positions is specifically DKC1-dependent. Furthermore, an analysis published data ^51–55^ demonstrated that a high stoichiometry of pseudouridine modification at 28S rRNA U4331 and U4966 is conserved across different cell lines (Figure S7B).

**Figure 7:**
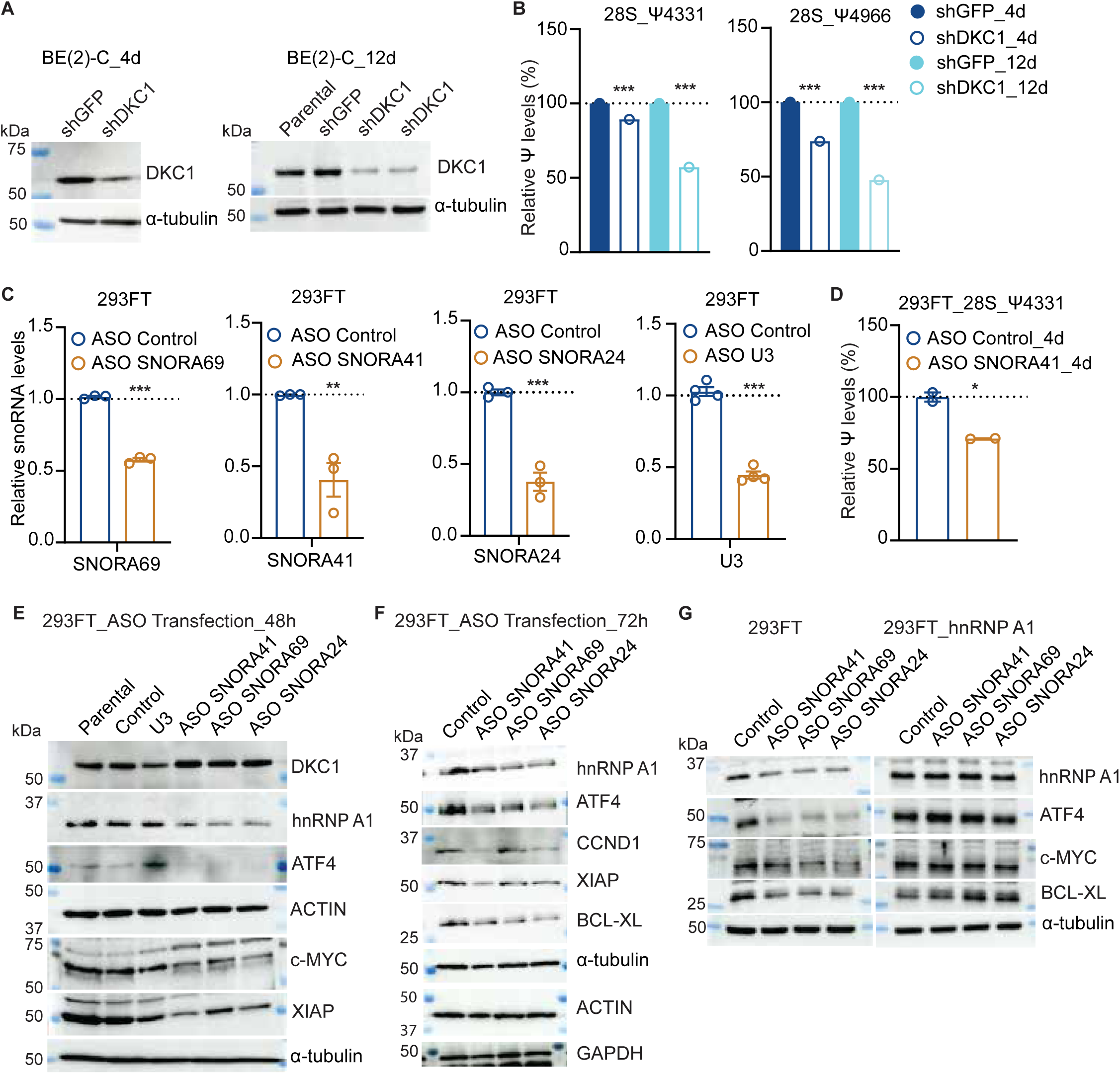
Functional Impact of Pseudouridylation at two 28S rRNA sites affected by DKC1. **(A)** Immunoblotting of DKC1 knockdown in BE(2)-C cells after 4 and 12 days of puromycin selection. **(B)** Relative pseudouridine levels at positions Ψ4331 and Ψ4966 in 28S rRNA from control and DKC1 knockdown BE(2)-C cells after 4 and 12 days of puromycin selection. Data were analyzed using the chi-square test. **(C)** qRT-PCR showing reduced expression of snoRNAs 69, 41, and 24 following transfection of 293FT cells with the indicated ASOs for 72 h. Data were analyzed using two-way ANOVA. **(D)** Relative pseudouridine levels at Ψ4331 in 28S rRNA from control or SNORA41 knockdown 293FT cells. Data were analyzed using the chi-square test. **(E-F)** Immunoblotting showing downregulation of hnRNP A1, ATF4, and IRES-dependent protein expression following knockdown of snoRNAs 41, 69, and 24 by ASOs at 48 h (**E**) and 72 h (**F**). **(G)** hnRNP A1 overexpression alleviated the inhibitory effect on ATF4 and IRES-dependent protein expression caused by ASO-mediated knockdown of snoRNAs 41, 69, and 24 in 293FT cells at 72 h. \**P* < 0.05, \*\**P* <0.01, \*\*\**P* <0.001.

Using the RNAsnoop tool ^56^, we identified potential box H/ACA snoRNAs associated with these Ψ sites: SNORA69 and SNORA41 for Ψ4331, and SNORA24 for Ψ4966 (Figure S7C). In addition, examination of published RNA interactome data revealed that DKC1 interacts with SNORA24 and SNORA41 in HepG2 cells ^46^ (Figure S7D). Further examination of published DKC1-RNA interactome data indicated that DKC1 binds to SNORA24 and SNORA69 ^57^ To investigate the functional significance of these snoRNAs, we utilized specific antisense oligonucleotides (ASOs) to reduce their expression, with an ASO targeting snoRNA U3 ^58^ serving as a control (Figure 7C). Furthermore, we examined whether the knockdown of specific snoRNAs would affect the expression of their corresponding host genes. SNORA41 is transcribed from EEF1B2, SNORA69 from RPL39, and SNORA24 from SNHG8 (Figure S7E). ASOs-mediated knockdown of these snoRNAs had no significant effect on mRNA expression levels of their host genes (Figure S7F).

As expected, for example, ASO-mediated SNORA41 knockdown decreased the pseudouridine level at Ψ4331 (Figure 7D), mimicking the effect of DKC1 knockdown. Importantly, knockdown of these snoRNAs decreased the protein expression of hnRNP A1 and ATF4, as well as IRES-dependent expression of MYC, XIAP, CCND1, and BCL-XL (Figures 7E and 7F). In contrast, knockdown of snoRNA U3 had no significant effect (Figure 7E). Furthermore, overexpression of hnRNP A1 alleviated the inhibitory effect of the ASOs on ATF4, MYC, and BCL-XL protein expression (Figure 7G), indicating that the effect of DKC1 on IRES-dependent translation is mediated through hnRNP A1.

Collectively, these findings suggest that DKC1-mediated rRNA pseudouridylation targets hnRNP A1 to drive IRES-dependent translation and ATF4 expression.

## Discussion

Our study identifies a DKC1-hnRNP A1 axis as critical for IRES-dependent translation and the activation of an ATF4-mediated transcriptional program essential for cancer cell survival and proliferation. Mechanistically, DKC1-mediated pseudouridylation of 28S rRNA enhances the translation of HNRNPA1 mRNA, leading to increased hnRNP A1 protein levels. hnRNP A1, in turn, binds to and stabilizes ATF4 mRNA, promoting ATF4 protein expression through IRES-mediated translation. This DKC1-hnRNP A1 axis is activated by the oncogene MYCN in neuroblastoma. We previously reported that MYCN upregulates ATF4 mRNA transcriptionally ^37^. By activating the DKC1-hnRNP A1 axis, MYCN further enhances ATF4 expression through a combination of transcriptional and translational mechanisms. ATF4 orchestrates the stress response to nutrient deprivation ^24–27,59^. Enhanced ATF4 expression provides a survival advantage to cancer cells under persistent stress conditions ^37,60–63^, which frequently arise due to the increased metabolic demands for biomass and energy production in rapidly proliferating tumors. Consequently, the DKC1-hnRNP A1 axis represents a specialized mechanism for the selective synthesis of proteins necessary to adapt to metabolic challenges, supporting cancer progression under stressful microenvironmental conditions.

Our study provides new mechanistic insights into how DKC1 regulates IRES-dependent translation. We show that DKC1-dependent pseudouridylation at two specific sites in 28S rRNA, Ψ4331 and Ψ4966, mediates the effect of DKC1 on IRES-dependent translation. It has been previously reported that pseudouridylation at these two sites are specifically and significantly reduced in ribosomes isolated from patients with familial dyskeratosis congenita ^51^. In agreement with the report, we found that DKC1 knockdown led to a time-dependent reduction in pseudouridine levels at these two sites. We identified three snoRNAs that potentially guide DKC1-mediated pseudouridylation at these two sites, SNORA41 and SNORA69 for Ψ4331, and SNORA24 for Ψ4966. Knockdown of these snoRNAs by ASOs fully recapitulated the effect of DKC1 knockdown on the expression of hnRNP A1 and ATF4, along with other proteins regulated by IRES-dependent translation. Our findings support the concept of specialized ribosomes, which can selectively translate subsets of mRNAs that contain unique regulatory elements, such as IRESs and upstream open reading frames (uORFs), enabling cells to fine-tune protein synthesis in response to diverse physiological demands ^64–66^.

In addition, our study identifies hnRNP A1, an ITAF ^44,48^, as a downstream effector of DKC1 in the regulation IRES-dependent translation. ITAFs promote IRES-dependent translation by facilitating ribosome recruitment to IRES in a cap-independent manner and/or by stabilizing the interaction of translation initiation factors with IRES ^48,67–69^. Several lines of evidence indicate that hnRNP A1 is a key downstream target of DKC1. At the functional level, hnRNP A1 knockdown or overexpression mimicked the effects of DKC1 knockdown or overexpression on the expression of ATF4 and its target genes, as well as on neuroblastoma cell and xenograft growth. At the molecular level, DKC1 is necessary for sustaining hnRNP A1 expression. Moreover, hnRNP A1 overexpression fully reversed the negative impact of DKC1 knockdown on IRES-dependent expression of BCL-XL, CCND1, MYC, and XIAP. Interestingly, we obtained evidence suggesting that hnRNP A1 interacts with and stabilizes DKC1, thereby creating a feed-forward loop that amplifies the activity of the DKC1-hnRNP A1 axis.

ATF4 is a master transcriptional regulator of the integrated stress response triggered by diverse stress signals, including amino acid deprivation and ER stress ^24–26^. It is well established that, under normal conditions, the translation of ATF4 mRNA is repressed due to inhibitory uORFs that prevent ribosomes from reaching the ATF4 coding region. During stress, eIF2α, a subunit of the translation initiation factor eIF2, is phosphorylated by stress-activated eIF2α kinases such as GCN2 (activated by the AAR) and PERK (activated by ER stress). Phosphorylation of eIF2α reduces active eIF2 levels, impairing the recruitment of the initiator Met-tRNA^Met^ to the 40S ribosomal subunit for translation initiation. This leads to a reduction in global mRNA translation ^26,70,71^. However, the decrease in initiating ribosomes alleviates repression at the uORFs in ATF4 mRNA, enabling efficient ATF4 translation. ^24–26^. Our findings reveal a mechanism for stress-induced ATF4 expression through IRES-dependent translation mediated by the DKC1-hnRNP A1 axis. A 2013 study identified an IRES element in the ATF4 V1 5’UTR, which is absent in the ATF4 V2 5’UTR ^50^. The ATF4 V1 IRES element promotes IRES-dependent translation in response to ER stress and eIF2α phosphorylation ^50^. It was speculated that the ATF4 V1 IRES element may interact with a cellular ITAF to stimulate its translation under stress ^50^. Our study identifies hnRNP A1 as the ITAF that drives ATF4 V1 expression during cellular stress. Co-transfection experiments demonstrated that hnRNP A1 induced much higher levels of ATF4 V1 protein expression compared to its effect on ATF4 V2 protein expression. Moreover, hnRNP A1 knockdown abolished ATF4 induction by both AAR and ER stress. Additionally, we showed that hnRNP A1 interacts with and stabilizes ATF4 mRNA. The role of hnRNP A1 in cellular stress responses is further supported by our observation that both AAR and ER stress induce hnRNP A1. Collectively, these findings provide compelling evidence and mechanistic insights into the critical role of IRES-dependent translation in stress-induced ATF4 protein expression.

Cancer cells often face challenging conditions, such as nutrient scarcity, hypoxia, and ER stress, due to their rapid proliferation and demanding microenvironment. To survive these conditions, cancer cells heavily rely on the translational machinery to selectively synthesize stress-related and survival proteins ^72^. Under these conditions, IRES-dependent translation becomes critical as canonical cap-dependent translation is downregulated. This alternative translation mechanism enables the selective production of proteins essential for cancer cell survival and adaptation. Not surprisingly, many stress-response mRNAs contain IRES elements in their 5’UTRs ^69,73,74^. Our findings reveal that MYCN activates the DKC1-hnRNP A1 axis, which enhances IRES-mediated translation and promotes ATF4-dependent upregulation of amino acid synthesis enzymes and transporters. This upregulation is vital for the growth and survival of MYCN-driven neuroblastoma. Thus, our study uncovers a mechanism by which an oncogene activates IRES-mediated translation to alleviate cellular stress, contributing to cancer development and progression.

## ACCESSION NUMBER

The BioProject accession number for the Illumina RNA-seq and nanopore RNA-seq data reported in this paper is PRJNA1196392.

## SUPPLEMENTAL INFORMATION

Supplemental Information includes 7 figures and 7 tables.

## ACKNOWLEDGMENTS

The authors extend their deepest gratitude to Dr. Sebla Kutluay at the University of Washington, St. Louis, for providing the pLHCX-hnRNP A1 plasmid and 293FT pLHCX-hnRNP A1 cells. They also thank Drs. Peter F. Stadler and Hakim Tafer at the University of Leipzig for their assistance with the RNAsnoop tool. Additionally, the authors express appreciation to Dr. Shuai Chen at the Sun Yat-sen University Cancer Center for sharing the DKC1 knockdown proteomics data and to Richard Kirkman and Dr. Suman Karki at the University of Alabama at Birmingham for their help with plasmid preparation and for providing the mouse anti-DKC1 antibody, respectively. Sunil Sudarshan is supported by VA Merit I01BX002930, R01CA200653, and DoD HT94252410558. The work was supported by R01CA190429 to Han-Fei Ding.

## AUTHOR CONTRIBUTIONS

A.G., M.B., and H-F.D. conceived the study and designed the experiments with contributions from J.D., M.P., and S.S. A.G. and M.B. performed the experiments, J.D. performed inducible DKC1 knockdown, co-immunoprecipitation, and co-transfection experiments, and M.P. performed polysome profiling. A.G., M.B., and H-F.D. analyzed data with contributions from J.D., M.P., and S.S. A.G., M.B., and H-F.D. wrote the paper. H-F.D. supervised and provided funding for the project. All authors read the manuscript and approved its contents.

## EXPERIMENTAL MODEL AND SUBJECT DETAILS

### Cell lines and cell culture

All the cell lines used in this study are listed in the Key Resources Table. Neuroblastoma cell lines BE(2)-C (CRL-2268) and SK-N-AS (CRL-2137) were acquired from ATCC, LA1-55n (06041203), from Sigma-Aldrich, and LA-N-6, SMS-KANR, and SMS-KCNR from the Children’s Oncology Group Cell Culture and Xenograft Repository, and SHEP1 from V.P. Opipari at the University of Michigan. 293FT (Thermo Fisher R70007), 293T Lenti-X (TaKaRa 632180), 293FT ATF4 KO ^77^, HeLa (ATCC CCL-2), SHEP1, SK-N-AS, and SHEP-MYCNER ^79^ cells were cultured in DMEM (HyClone SH30022), while BE(2)-C was maintained in DMEM/Ham’s F-12 (1:1) (HyClone SH30023). All other cell lines were cultured in RPMI 1640 (HyClone SH30027). The 293FT pLHCX hnRNP A1 (NM_002136.4) cell line ^76^ was a gift from Sebla B. Kutluay at Washington University School of Medicine. All media were supplemented with 10% FBS (Atlanta Biologicals; S11050). All cell lines were authenticated through short tandem repeat (STR) profiling. After authentication, extensive frozen stocks were created to prevent cross-contamination. Cell lines were utilized within 10 passages post-thawing and were regularly screened for Mycoplasma contamination every three months using the LookOut Mycoplasma PCR Kit (Sigma-Aldrich) and DAPI staining. Phase-contrast microscopy images of cells were captured using the EVOS M5000 Imaging System (Invitrogen). Cell proliferation was assessed using a Trypan blue assay.

The small-molecule DKC1 inhibitor Pyrazofurin was purchased from Sigma-Aldrich (Cat# SML1502), dissolved in water, and stored at −80°C. Cells were treated with various concentrations of Pyrazofurin for 3 days before being collected for cell counting, qRT-PCR, and immunoblot analyses

For cellular stress response experiments, cells were treated for 5 hours with either vehicle (H₂O for HisOH control or DMSO for tunicamycin control), 5 mM Histidinol (HisOH, Sigma-Aldrich H6647), or 2 µg/mL tunicamycin (Sigma-Aldrich T7765), and the collected for immunoblot analysis.

### Animal experiments

For xenograft studies, we used 6-week-old male and female NOD.SCID mice (NOD.Cg-Prkdcscid/J, 001303) obtained from the Jackson Laboratory. BE(2)-C cells expressing shGFP (control), shDKC1 738, shDKC1 326, or shHNRNPA1 586 were suspended in 100 µl of Hanks’ Balanced Salt Solution (Thermo Fisher 14170112) and injected subcutaneously into both flanks of the mice (two sites per mouse) at approximately 4 × 10 cells per site. Tumor volume was measured every other day with a digital caliper and calculated using the formula V = (L × W²)/2. Mice were euthanized when tumors reached ∼1.0 cm in any diameter. All animal procedures were conducted at the University of Alabama at Birmingham and approved by the Institutional Animal Care and Use Committee (IACUC).

### Patient data

The neuroblastoma patient RNA-seq data used in this study were collected as described previously ^63,80^. Survival and gene expression correlation analyses were conducted using the R2: Genomics Analysis and Visualization Platform ^80^ (https://hgserver1.amc.nl/cgi-bin/r2/main.cgi). The resulting figures and p-values were subsequently downloaded for further analysis.

### Overexpression and knockdown

Lentiviral constructs for overexpressing human DKC1 (pLenti6.3/V5-DEST, HsCD00943802) were obtained from DNASU. Human DKC1 shRNA oligonucleotide sequences shDKC1 738 (TRCN0000039738), shDKC1 326 (TRCN0000010326), and shDKC1 740 (TRCN0000039740) were obtained from the Broad Institute (https://portals.broadinstitute.org/gpp/public/gene/search), amplified by PCR, subcloned into the pLKO.1 puro vector, and validated by sequencing. In addition, shDKC1 sequences (TRCN0000039743 and TRCN0000332892) were cloned into Tet-pLKO-puro (Addgene 21915) for inducible expression in the presence of 0.5 µg/mL doxycycline.

The lentiviral construct pLHCX-hnRNP A1 for overexpressing human hnRNP A1 variant 1 (NM_002136.4) was kindly provided by Sebla B. Kutluay (Washington University School of Medicine, St. Louis, MO). The DNA sequence coding for hnRNP A1 variant 2 (NM_031157.4) was obtained from DNASU (HNRNPA1 in pANT7_cGST, HsCD00785011), amplified by PCR, subcloned into pCDH-CMV-MCS-EF1-puro (SBI System Biosciences CD510B-1), and validated through sequencing. Human HNRNPA1 shRNA sequences—shHNRNPA1 566 (TRCN0000147566), shHNRNPA1 582 (TRCN0000006582), shHNRNPA1 585 (TRCN0000006585), shHNRNPA1 586 (TRCN0000006586), and shHNRNPA1 048 (TRCN0000148048)—were obtained from the Broad Institute, amplified by PCR, subcloned into pLKO.1 puro, and verified by sequencing.

Lentiviruses were produced in 293T Lenti-X cells using packaging plasmids pLP1 (Addgene_22614), pLP2, and pLP/VSVG (Thermo Fisher Scientific K497500). Lentiviral infection of cells was conducted according to standard procedures.

Co-transfection of 293FT ATF4 KO cells was conducted with pGenLenti-ATF4 (V1, NM_001675, GenScript) ^63^ or pCW57.1-ATF4 (V2, NM_182810) ^77^, alongside DKC1- and/or HNRNPA1-expressing plasmids. Cells were collected 48 hours after transfection for immunoblot analysis.

### Quantitative RT-PCR

Total RNA was extracted from cells using TRIzol reagent (Thermo Fisher, 15596026). Reverse transcription was performed with the iScript Advanced cDNA Synthesis Kit (Bio-Rad, 172-5038). Quantitative RT-PCR was carried out using a 2X SYBR Green qPCR Master Mix (Bimake, B21203) on an iQ5 real-time PCR system (Bio-Rad) with gene-specific primers (Supplementary Table 6). mRNA levels were normalized to β2-microglobulin (B2M). Primer specificity was confirmed through melting curve analysis after qRT-PCR, ensuring that each primer pair produced a single, distinct amplification peak.

To measure mRNA half-life, BE(2)-C shGFP and shHNRNPA1 585 cells were treated with 5 µg/mL Actinomycin D (Sigma-Aldrich A9415) for various times (0, 0.5, 1, 2, and 4 hours), followed by total RNA extraction and qRT-PCR analysis.

### Immunoblotting

Cell lysates were prepared using a standard SDS sample buffer, and protein concentrations were measured using the Bio-Rad Protein Assay Kit II (5000002). Proteins (15–35 µg) were resolved by SDS-PAGE and transferred onto nitrocellulose membranes. The membranes were then blocked for 60 minutes at room temperature with gentle shaking in 5% nonfat milk prepared in Tris-buffered saline containing 0.1% Tween 20 (TBS-T). Immunoblotting was performed using the following primary antibodies: rabbit anti-ASNS (1:1000, Proteintech 14681-1-AP), rabbit anti-ATF4 (D4B8, 1:1000, Cell Signaling 11815), rabbit anti-BCL-XL (54H6, 1:1000, Cell Signaling 2764S), mouse anti-CCND1 (DCS-6, 1:200, Santa Cruz sc-20044), rabbit anti-DKC1 (1:1000, Bethyl Laboratories A302-591A), mouse anti-DKC1 (1:500, Santa Cruz sc-373956), rabbit anti-GAPDH (FL-335, 1:1000, Santa Cruz sc-25778, RRID), rabbit anti-hnRNP A1 (1:15,000, Proteintech 11176-1-AP), mouse anti-hnRNP A1 (4B10, 1:2000, Santa Cruz sc-32301), rabbit anti-MTHFD2 (1:1500, Proteintech 12270-1-AP), mouse anti-MYCN (B8.4.B, 1:400, Santa Cruz sc-53993), mouse anti-p27KIP1 (1:200, Santa Cruz sc-1641), rabbit anti-PHGDH (1:300, Sigma-Aldrich HPA021241), rabbit anti-PSAT1 (1:3000, Novus 21020002), rabbit anti-SHMT2 (1:1000, Millipore Sigma HPA020549), rabbit anti-SLC7A5/LAT1 (1:1000, Cell Signaling 5347), rabbit anti-SLC7A11/xCT (D2M7A, 1:1000, Cell Signaling 12691), rabbit anti-β-actin (1:2000, Thermo Fisher Scientific MA5-15739, RRID), mouse anti-α-tubulin (B-5-1-2, 1:5000, Sigma-Aldrich T5168, RRID), and rabbit anti-XIAP (1:1000, Cell Signaling 2042).

The secondary antibodies used were horseradish peroxidase (HRP)-conjugated goat anti-mouse (Jackson ImmunoResearch 115-035-146) and goat anti-rabbit IgG (Jackson ImmunoResearch 111-035-046). Immunoblots were visualized using the Clarity Western ECL Substrate Kit (Bio-Rad 1705061) and quantified with the Amersham ImageQuant 800 (Cytiva) or ImageJ software (version 1.53k).

### Co-immunoprecipitation

Nuclear extracts were prepared from BE(2)-C cells as previously described ^81^ and incubated overnight at 4°C with Protein A magnetic beads (Bio-Rad SureBeads 161-4011) coated with 2 µg of rabbit IgG (Santa Cruz sc-2027), rabbit anti-DKC1 (Bethyl Laboratories A302-591A), or rabbit anti-hnRNP A1 (Proteintech 11176-1-AP). The beads were collected using a magnetic separator, washed with extraction buffer, and suspended in standard SDS sample buffer for immunoblot analysis.

### RNA Immunoprecipitation

RNA immunoprecipitation was performed as described previously ^82^, with minor modifications. Briefly, BE(2)-C cells were washed with ice-cold PBS and lysed in a lysis buffer (20 mM Tris-HCl, pH 7.5, 100mM KCl, 5 mM MgCl2, and 0.5% NP-40) for 10 minutes on ice. The lysate was then centrifuged at 10,000 ×g for 15 minutes at 4°C. The supernatant was collected and incubated overnight at 4°C with Protein A magnetic beads (Bio-Rad SureBeads 161-4011) coated with 2 µg of mouse IgG (Santa Cruz sc-2025 or mouse anti-hnRNP (4B10, Santa Cruz sc-32301). The beads were collected using a magnetic separator, washed with a buffer (50 mM Tris-HCl, pH 7.5, 150 mM NaCl, 1 mM MgCl₂, and 0.05% NP-40), and treated with 20 U of RNase-free DNase I for 15 minutes at 37°C. A portion of the sample was collected for immunoblotting, while the remaining sample was treated with 0.1% SDS and 0.5 mg/mL Proteinase K for 15 minutes at 55°C to remove proteins. RNA was then extracted using TRIzol and analyzed via qRT-PCR.

### Metabolomics

BE(2)-C_teton and BE(2)-C_teton-shDKC1-92 cells were cultured in the presence of 0.5 μg/mL doxycycline for 6 days, while BE(2)-C cells were treated with 1 µM pyrazofurin for 3 days. Cells were then collected by scraping and centrifugation, washed once with ice-cold PBS, snap-frozen in liquid nitrogen, and stored at −80°C for metabolomics analysis. Six biological replicates (∼5×10 cells/sample) were analyzed for each group. Untargeted metabolomics profiling was performed at the NIH West Coast Metabolomics Center at the University of California, Davis. Pathway analysis of metabolites was performed using MetaboAnalyst ^34^.

### Antisense oligonucleotides (ASO’s) transfection

ASO transfection was performed according to a published protocol ^83^. The ASO sequences are listed in Supplementary Table 7. Briefly, 293FT cells were seeded in a 6-well plate at a density optimized to achieve approximately 70% confluency within 24-48 hours. The ASO-Polyethylenimine (PEI) complex was prepared by mixing 5 µL of 50 µM ASO with 3.5 µL of PEI (1 mg/mL) in 91.5 µL of sterile 0.9% saline solution. The mixture was thoroughly pipetted to ensure homogenization and then incubated at room temperature for 15 minutes.

Prior to transfection, the cells were washed once with PBS, and 900 µL of fresh medium was added to each well and the plate was incubated for 15 minutes. The ASO-PEI complex was then added dropwise to the cells, bringing the final concentration of ASO in the medium to 250 nM. The plate was gently shaken to ensure even distribution of the transfection solution and incubated at 37°C for 5-6 hours. After the incubation, the medium was carefully removed, and 2 mL of fresh medium was added to each well. The cells were cultured for 48-72 hours to ensure optimal ASO uptake and then were collected for qRT-PCR, immunoblotting, and Nanopore direct RNA sequencing.

### Poly-A RNA sequencing

Total RNA was prepared from BE(2)-C control, BE(2)-C shDKC1 738, and BE(2)-C treated with 1 µM Pyrazofurin for 72 hours. Poly-A RNA was extracted from high-quality total RNA (RIN > 9.0). The poly-A RNA samples from the control and Pyrazofurin-treated BE(2)-C cells was paired-end sequenced using the Illumina HiSeq 4000 platform by Azenta Life Sciences. Following base calling by Illumina bcl2fastq (v2.18.0.12), FASTQ files were aligned to the Homo_sapiens.GRCh38.cdna.all reference assembly using the Kallisto tool (https://github.com/pachterlab/kallisto). Transcript abundance was quantified using the edgeR package (https://bioconductor.org/packages/release/bioc/html/edgeR.html), and differential expression analysis was performed with the limma-voom package (https://bioconductor.org/packages/release/bioc/html/limma.html).

For Nanopore Direct RNA sequencing, BE(2)-C control and BE(2)-C shDKC1 738 mRNA libraries were prepared as described previously ^84^. Raw fast5 files were basecalled using Guppy (v6.4.2), and the resulting FASTQ files were aligned to the reference sequence (GRCh38.p13) using minimap2 ^85^ (https://github.com/lh3/minimap2) and Bambu ^86^ (https://github.com/GoekeLab/bambu) was used to estimates transcripts abundance for differential gene expression analysis. Gene Ontology (GO) analysis and Gene Set Enrichment Analysis (GSEA) were performed using enrichplot R tool (https://bioconductor.org/packages/release/bioc/html/enrichplot.html).

### rRNA sequencing

Total RNA was isolated from BE(2)-C control, BE(2)-C shDKC1 738, BE(2)-C shPUS7-33, 293FT ASO control, and 293FT ASO SNORA41 cells. rRNA was enriched from total RNA by depleting poly-A RNA using a Poly(A) mRNA Magnetic Isolation Module (NEB E7490S). A poly-A tail was then added to rRNA using a poly(A) Tailing Kit (NEB M0276). The direct rRNA sequencing library was prepared according to the standard protocol and loaded into a Flongle flow cell for sequencing using MinKnow (version 23). Each flow cell was sequenced for 24 hours. Raw fast5 files were converted to pod5 files for basecalling using Dorado v8 (https://github.com/nanoporetech/dorado). The resulting FASTQ files were aligned to a human non-coding RNA reference sequences, and pseudouridine sites were identified based on significant changes in U-to-C basecalling errors ^84^.

### Polysome profiling

Polysome analysis was performed as described previously ^87^, with minor modifications. BE(2)-C cells expressing shGFP and shDKC1 326 were treated with cycloheximide (Sigma-Aldrich, Cat# C4859) at 0.1 mg/mL for 3 minutes, washed with PBS containing 0.1 mg/mL cycloheximide, and lysed on ice for 15 minutes using polysome extraction buffer (15 mM Tris-Cl, pH 7.4, 15 mM MgCl₂, 0.3 M NaCl, 0.1 mg/mL cycloheximide, 1 mg/mL heparin, and 1% Triton X-100). The lysates were centrifuged at 13,200 xg for 10 minutes, and 0.5 mL of the supernatant was layered onto 10%-50% sucrose gradients in polysome extraction buffer (without Triton X-100). The gradients were centrifuged at 41,000 xg for 2 hours in an SW41 Ti rotor (Beckman, Cat# BR-8101) at 4°C. Fractions were collected using a Gradient Station IP system (BioComp) into open-top polyclear centrifuge tubes (Seton Scientific, Cat# 7030) and stored at −80°C. Total RNA was extracted from 0.5 mL of each polysome fraction using Acid Phenol: Chloroform (125:24.1, pH 4.5 ± 0.2, Ambion, Cat# AM9720). The RNA was then precipitated with 70% ethanol and analyzed by qRT-PCR. Data were normalized to GAPDH.

### snoRNA-rRNA Interaction

For the snoRNA-rRNA interaction analysis, we utilized the RNA snoop tool as previously described ^56^. H/ACA snoRNA sequences were obtained from the snoDB database ^83^. Prior to running RNA snoop to accurately identify snoRNAs associated with DKC1-dependent pseudouridine sites, the snoRNA sequences were preprocessed by removing the H box (ANANNA) and ACA box, as described previously ^56^. This preprocessing step resulted in a database of 363 unique snoRNA sequences. Subsequently, we used the 28S rRNA as the input RNA to identify snoRNAs targeting pseudouridine sites at 28S_U4331 and 28S_U4966.

### Venn diagram analysis

Venn diagram analysis was conducted using three distinct datasets: 1) Genes positively correlated with DKC1 expression - This dataset was derived from the SEQC neuroblastoma patient dataset ^80^, applying a correlation threshold of R > 0.7 and a p-value threshold of < 1.143E-72; 2) Genes downregulated following DKC1 inhibition - This dataset included genes with a fold change of ≥ −3.0 and an adjusted p-value of < 0.01; and 3) Known ITAF factors - This dataset was obtained from the IRESite database (http://www.iresite.org/). The overlap among these three gene sets was visualized using the R package VennDiagram ^88^.

### Statistical Analysis

Quantitative data are expressed as mean ± SEM and were analyzed for statistical significance using an unpaired, two-tailed Student’s t-test or ANOVA (one-way or two-way). Dose-response curves for the DKC1 inhibitor were fitted using the “log(inhibitor) vs. response (three parameters)” model. Pseudouridine modification data was analyzed by using the Chi-square test. For animal studies, the log-rank test was applied to assess mouse survival at the end of the experiment. Unless otherwise specified, all statistical analyses were performed using GraphPad Prism 10.2.3 for Windows. Statistical significance was assigned as *P < 0.05; **P < 0.01; ***P < 0.001.

### Data availability

The BioProject accession number for the Illumina RNA-seq and nanopore RNA-seq data reported in this paper is PRJNA1196392. The patient data analyzed in this study were obtained from R2 Genomics Analysis and Visualization Platform (https://hgserver1.amc.nl/cgi-bin/r2/main.cgi). All other raw data are available upon request from the corresponding authors.

---

**Figure S1.**
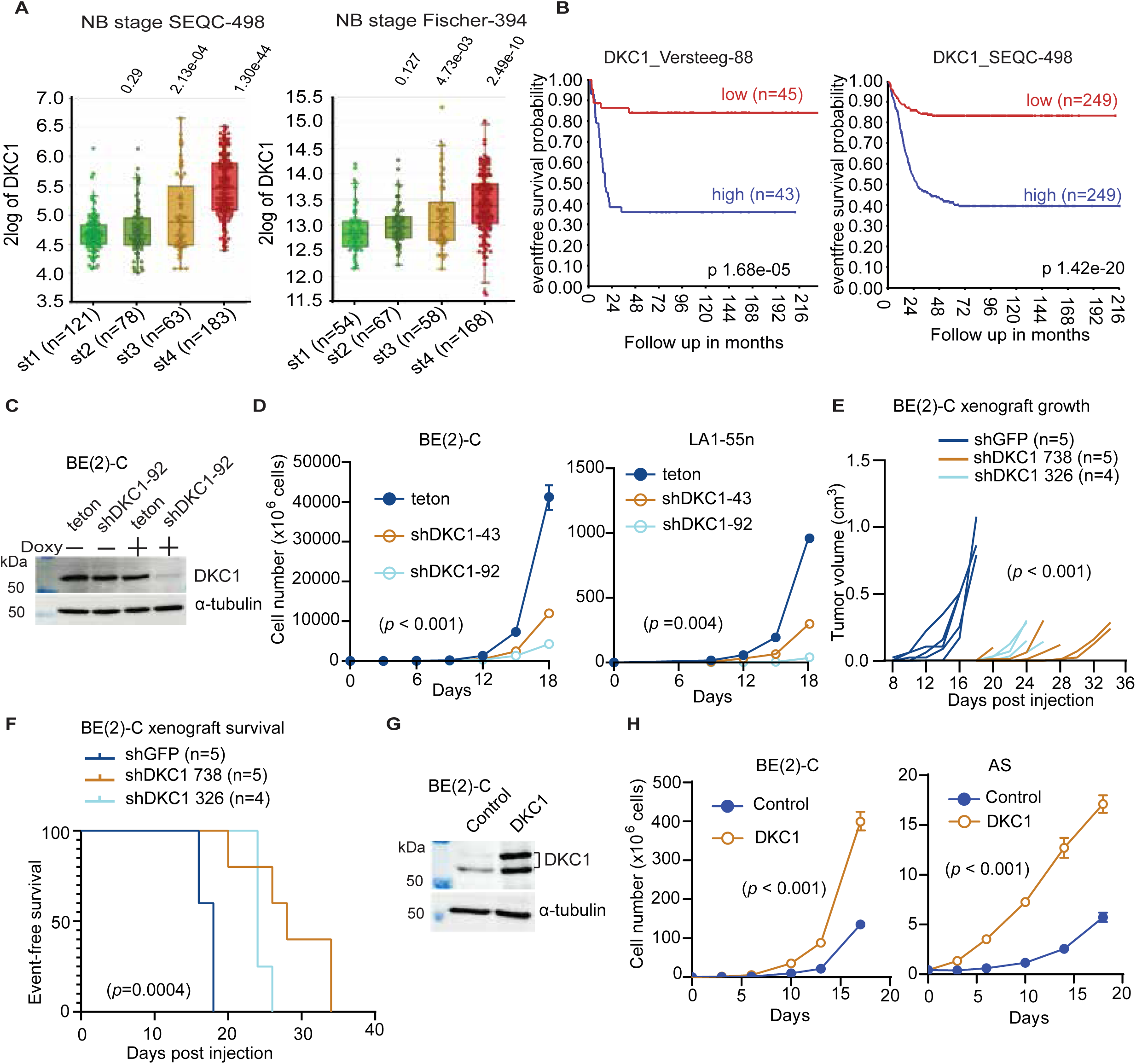
High DKC1 expression is associated with poor prognosis and is required for the proliferation and tumorigenicity of neuroblastoma cells. **(A)** Box plots showing increased DKC1 mRNA expression in neuroblastoma tumors at advanced stages, based on datasets from the SEQC cohort (n = 498) and the Fischer cohort (n = 394). *P* values were determined by one-way ANOVA. **(B)** Kaplan-Meier survival curves for two independent cohorts of neuroblastoma patients based on DKC1 mRNA expression, with indicated log-rank P values. High DKC1 expression is associated with poor prognosis. **(C)** Immunoblotting showing inducible DKC1 knockdown in the presence of doxycycline (Doxy+, 0.5µg/mL for 6 days). **(D)** DKC1 knockdown (Doxy+) reduced the proliferation of neuroblastoma BE(2)-C and LA1-55n cells. **(E-F)** DKC1 knockdown impeded BE(2)-C xenograft growth **(E)** and prolonged event-free survival **(F)** of xenograft-bearing NOD.SCID mice. Log-rank test P value is indicated. **(G)** Immunoblotting showing DKC1 overexpression. **(H)** DKC1 overexpression promoted the proliferation of neuroblastoma BE(2)-C and SK-N-AS cells. Cell growth data (**D** and **H**) are presented as mean ± SEM (n = 4) were analyzed using two-way ANOVA.

**Figure S2.**
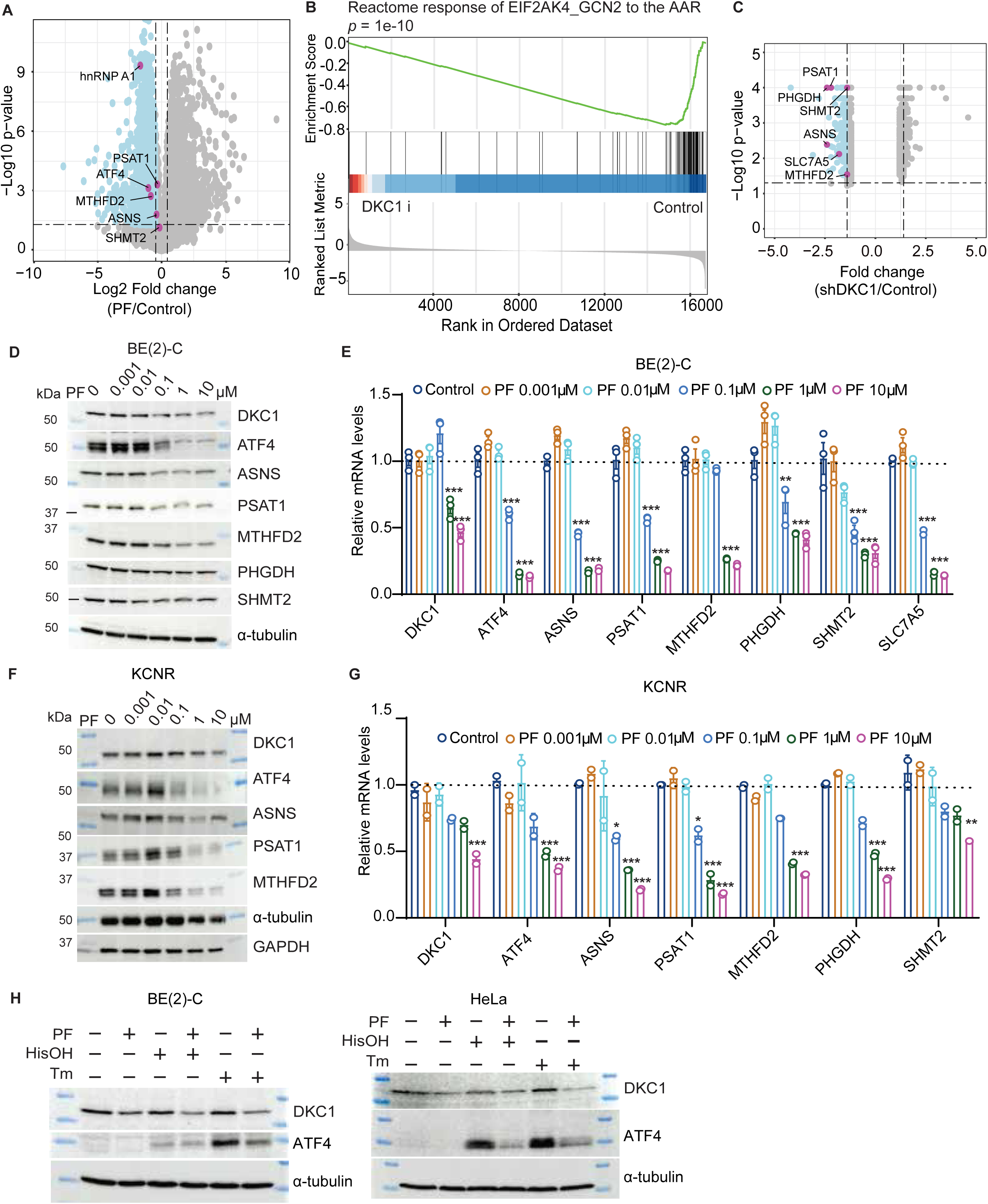
DKC1 is essential to sustain the expression of genes involved in amino acid metabolism and cellular stress responses. **(A)** Volcano plot of RNA-seq data showing downregulation of genes involved in amino acid metabolism after DKC1 inhibition for 72 h in BE(2)-C cells. **(B)** GSEA of RNA-seq data showing downregulation of genes involved in the EIF2AK4_GCN2 response to amino acid deficiency following DKC1 inhibition. **(C)** Volcano plot of proteomics data showing downregulation of proteins involved in amino acid metabolism in the colorectal adenocarcinoma cell line DLD-1 following DKC1 inhibition. **(D-G)** qRT-PCR **(D, F)** and immunoblotting **(E, G)** showing downregulation of mRNA and protein expression of representative genes involved in amino acid metabolism following DKC1 inhibition in neuroblastoma BE(2)-C and KCNR cells. qRT-PCR data are presented as mean ± SEM (n = 3) and analyzed using two-way ANOVA. **(H)** Immunoblotting showing DKC1 inhibition abrogated stress-induced upregulation of ATF4 in BE(2)-C and HeLa cells. Cells were treated without or with the DKC1 inhibitor pyrazofurin 1µM for 48 h, followed by treatment with 5µM Histidinol (HisOH, AAR inducer) or 2µg/mL tunicamycin (Tm, ER stress inducer) for 6 h. \*\**P* < 0.01, \*\*\**P* <0.001.

**Figure S3.**
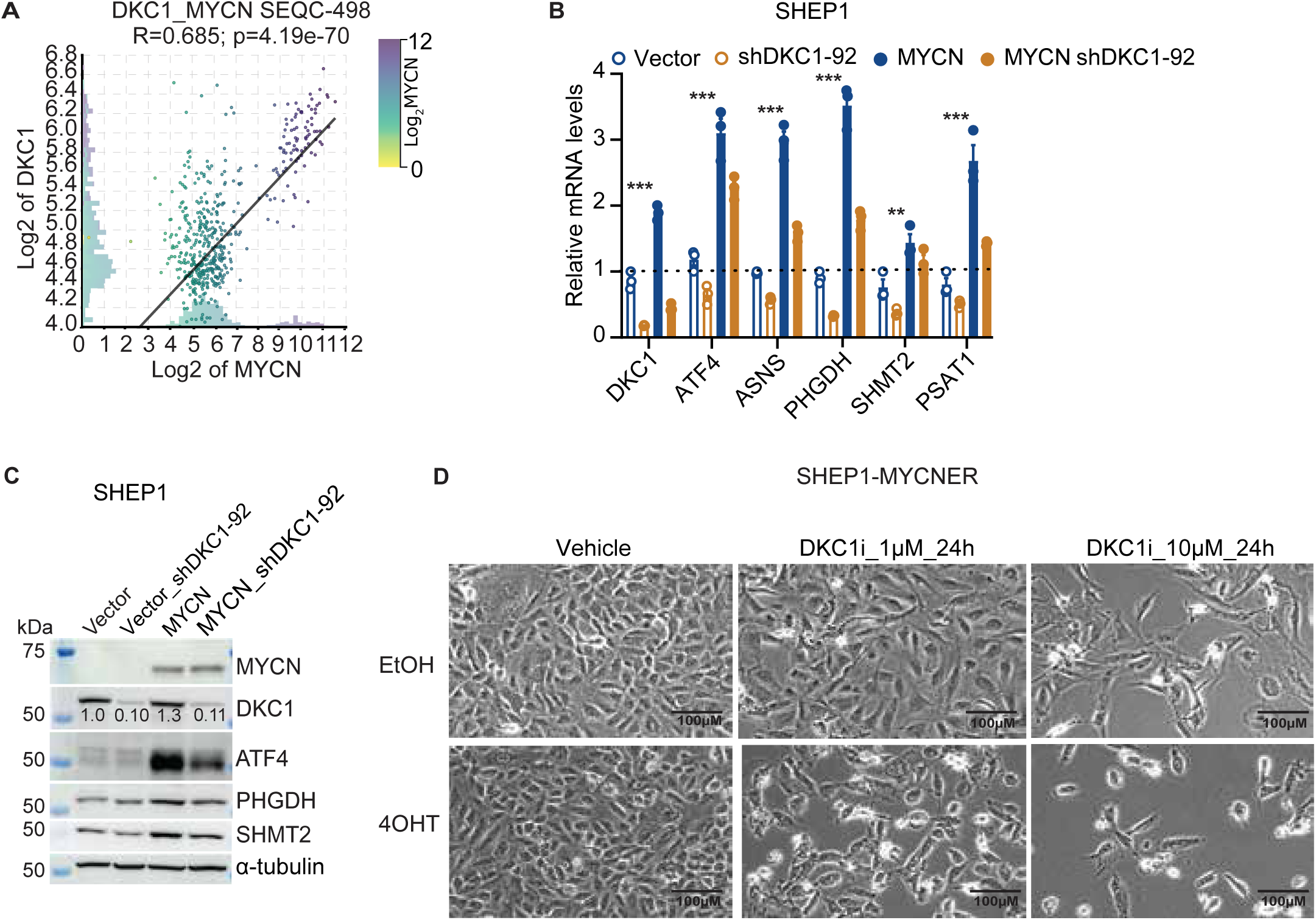
DKC1 is a downstream effector of MYCN. **(A)** Positive correlation in mRNA expression between DKC1 and MYCN in primary neuroblastoma tumors (the SEQC cohort, n = 498), R (correlation) and p values are indicated. **(B-C)** qRT-PCR (**B**) and Immunoblotting **(C)** showing DKC1 knockdown abrogated MYCN upregulation of genes for amino acid synthesis in SHEP1 cells. qRT-PCR data are presented as mean ± SEM (n = 3) and analyzed using two-way ANOVA. **(D)** Phase contrast imaging showing MYCN activation by 4OHT sensitized cells to DKC1 inhibition. \*\**P* <0.01, \*\*\**P* <0.001.

**Figure S4.**
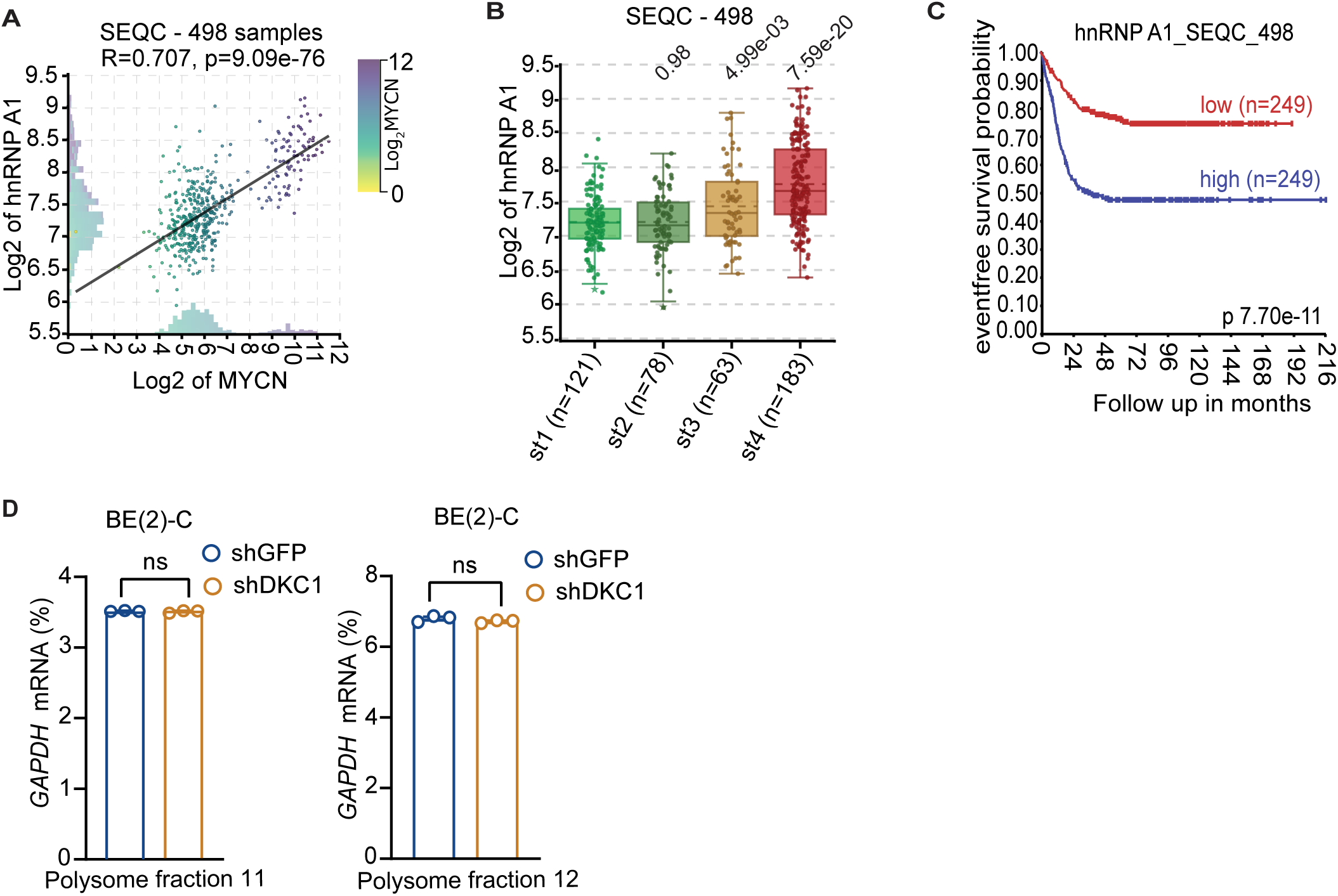
High HNRNPA1 mRNA expression is associated with poor prognosis in neuroblastoma patients. **(A)** Positive correlation in mRNA expression between HNRNPA1 and MYCN in primary neuroblastoma tumors (the SEQC cohort, n = 498), R (correlation) and p values are indicated. **(B)** Box plots showing increased HNRNPA1 mRNA expression in neuroblastoma tumors at advanced stages, based on the SEQC dataset (n = 498). **(C)** Kaplan-Meier survival curves for the SEQC cohort of neuroblastoma patients based on HNRNPA1 mRNA expression with indicated log-rank *P* value. High HNRNPA1 expression is associated with poor prognosis. **(D)** qRT-PCR analysis of polysome fractions showing DKC1 knockdown had no effect on polysome-associated GAPDH mRNA levels. Data represent two independent experiments and are presented as the mean ± SEM of 3 technical replicates. ns, not significant.

**Figure S5.**
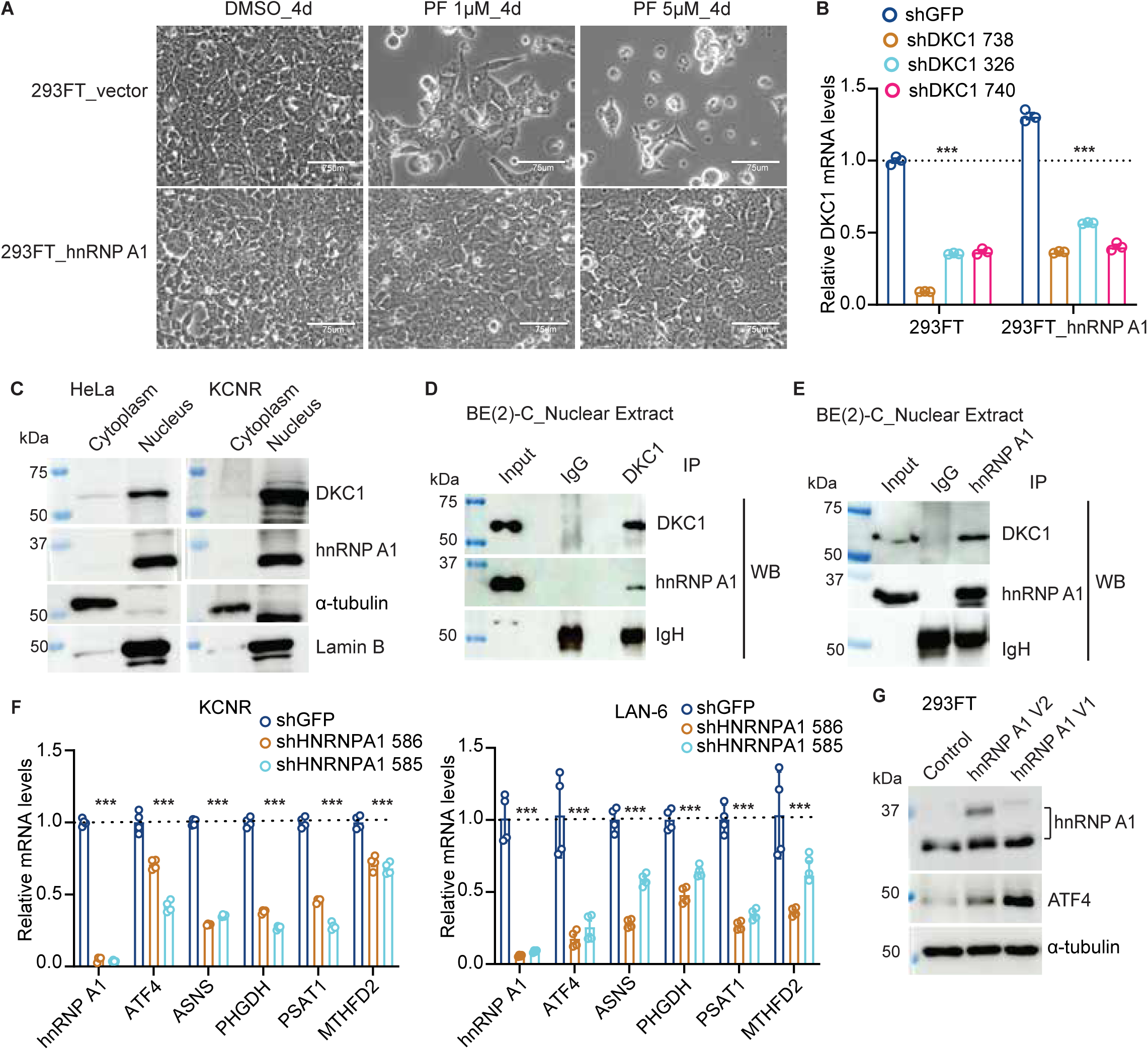
hnRNP A1 interacts with DKC1 and sustains the expression of ATF4 and genes involved in amino acid metabolism. **(A)** Phase contrast imaging showing hnRNP A1 overexpression alleviated the inhibitory effect of DKC1 inhibition on 293FT cell survival. **(B)** qRT-PCR showing hnRNP A1 overexpression had no significant effect on shRNA-mediated knockdown of DKC1 mRNA expression in 293FT cells. data are presented as mean ± SEM (n = 3) and analyzed using two-way ANOVA. **(C)** Co-localization of hnRNP A1 and DKC1 in the nucleus of HeLa and neuroblastoma SMS-KCNR cells, α-tubulin and Lamin B were used as cytoplastic and nuclear markers, respectively. **(D-E)** Immunoblot analysis of co-immunoprecipitation of hnRNP A1 and DKC1 in nuclear extracts from BE(2)-C cells. **(F)** qRT-PCR showing downregulation of representative genes involved in amino acid metabolism following hnRNP A1 knockdown in SMS-KCNR and LAN-6 cells. qRT-PCR data are presented as mean ± SEM (n = 4) and analyzed using two-way ANOVA. **(G)** Immunoblotting showing upregulation of ATF4 expression following overexpression of hnRNP A1 V1 and V2. \*\*\**P* < 0.001.

**Figure S6.**
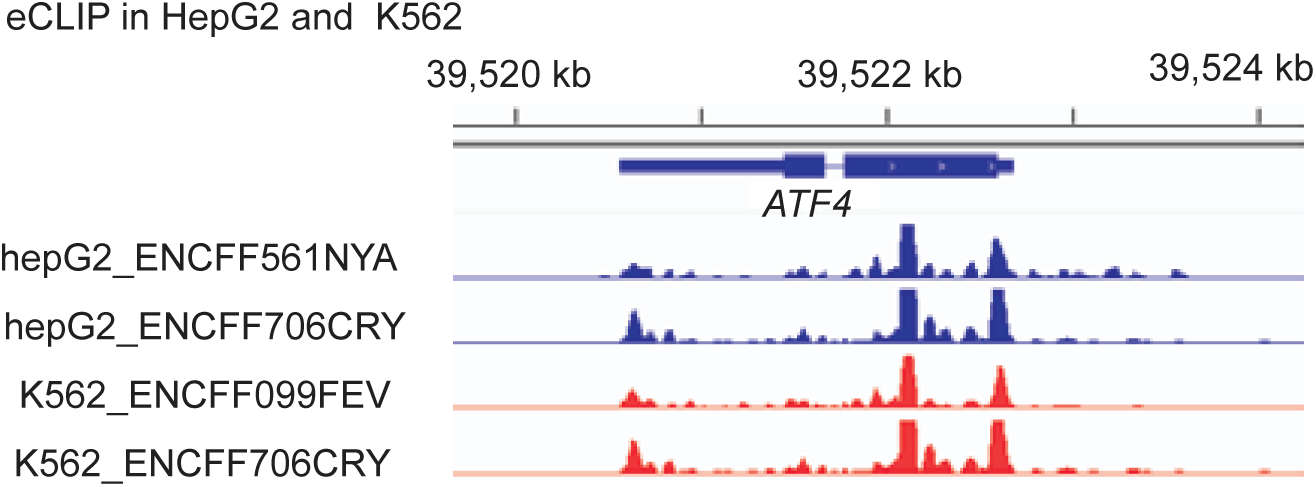
hnRNP A1 binds to ATF4 mRNA. Integrative genomics viewer (IGV) of hnRNP A1 eCLIP analysis showing hnRNP A1 binding to Atf4 mRNA, with the indicated peaks representing hnRNP A1-bound regions.

**Figure S7.**
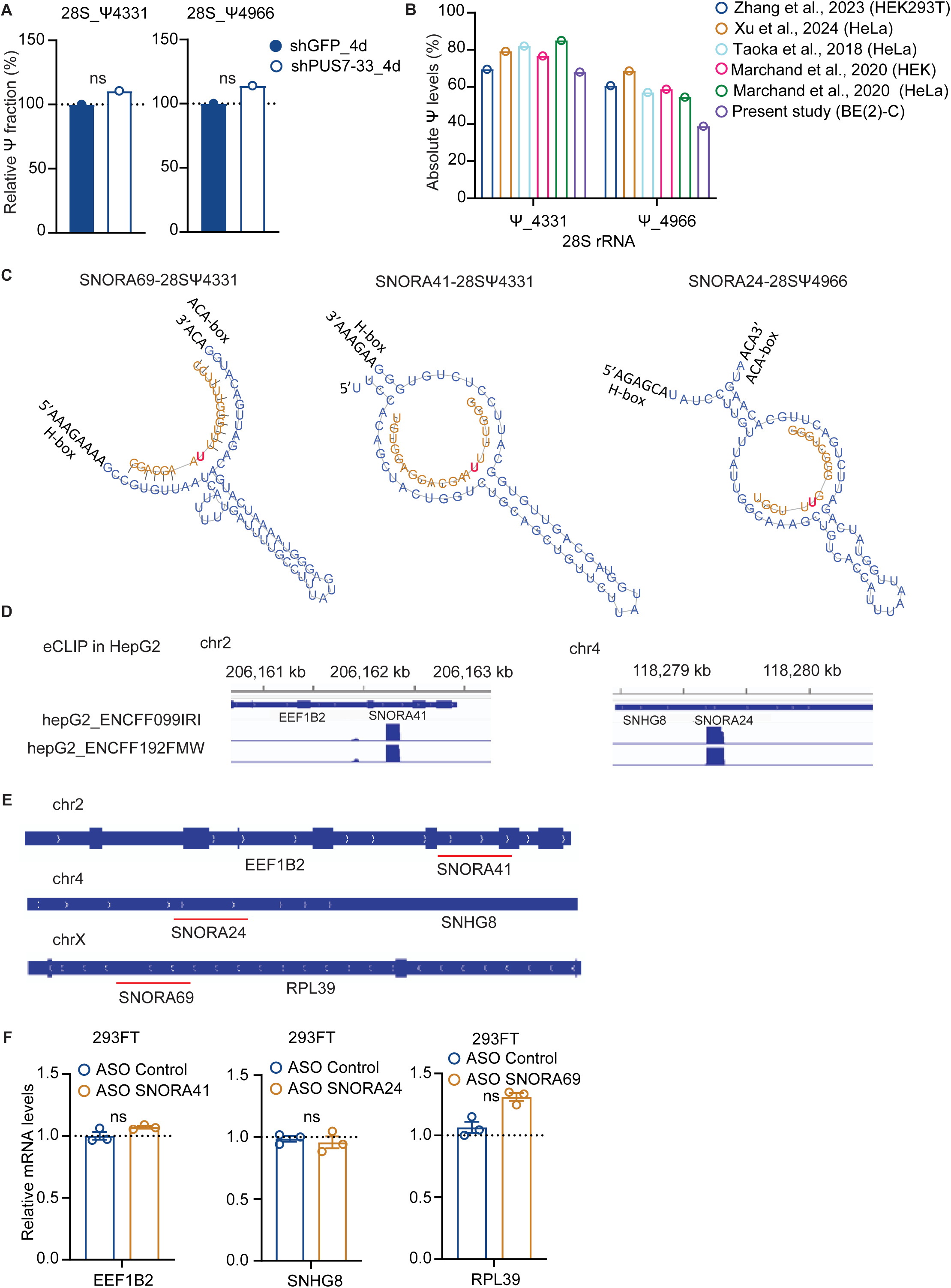
DKC1-mediated pseudouridylation at 28S rRNA positions Ψ4331 and Ψ4966. **(A)** Relative pseudouridine levels at 28S rRNA Ψ4331 and Ψ4966 in BE(2)-C cells following PUS7 knockdown. **(B)** Analysis of absolute pseudouridine levels at 28S rRNA Ψ4331 and Ψ4966 across various cell lines using published data. **(C)** The hairpin-ψ pocket interaction of human box H/ACA SNORA69, SNORA41, and SNORA24 guide RNAs with 28S rRNA at Ψ4331 and Ψ4966. The target rRNA sequences are represented in brown, while the corresponding snoRNA sequences are shown in blue. The pseudouridylation sites are highlighted in red. **(D)** IGV of DKC1 eCLIP analysis showing DKC1 binding to SNORA24 and SNORA41 in HepG2 cells, with the indicated peaks representing DKC1 bound regions. **(E)** IGV of EEF1B2, SNHG8 and RPL39 transcripts with their corresponding H/ACA snoRNAs, SNORA41, SNORA24, and SNORA69, highlighted in red underlined. (**F**) qRT-PCR showing relative mRNA expression of host genes EEF1B2, SNHG8, and RPL39 following ASO-mediated knockdown of SNORA41, SNORA24, and SNORA69, respectively. ns, not significant.

## References

1. Heiss, N.S., Knight, S.W., Vulliamy, T.J., Klauck, S.M., Wiemann, S., Mason, P.J., Poustka, A., and Dokal, I. (1998). X-linked dyskeratosis congenita is caused by mutations in a highly conserved gene with putative nucleolar functions. Nat Genet 19, 32–38. 10.1038/ng0598-32.

2. Knight, S., Vulliamy, T., Copplestone, A., Gluckman, E., Mason, P., and Dokal, I. (1998). Dyskeratosis Congenita (DC) Registry: identification of new features of DC. British Journal of Haematology 103, 990–996. 10.1046/j.1365-2141.1998.01103.x.

3. Dokal, I. (2000). Dyskeratosis congenita in all its forms. British Journal of Haematology 110, 768–779. 10.1046/j.1365-2141.2000.02109.x.

4. Ruggero, D., Grisendi, S., Piazza, F., Rego, E., Mari, F., Rao, P.H., Cordon-Cardo, C., and Pandolfi, P.P. (2003). Dyskeratosis Congenita and Cancer in Mice Deficient in Ribosomal RNA Modification. Science 299, 259–262. doi:10.1126/science.1079447.

5. Koonin, E.V. (1996). Pseudouridine Synthases: Four Families of Enzymes Containing a Putative Uridine-Binding Motif Also Conserved in dUTPases and dCTP Deaminases. Nucleic Acids Research 24, 2411–2415. 10.1093/nar/24.12.2411.

6. Hamma, T., and Ferre-D’Amare, A.R. (2006). Pseudouridine synthases. Chem Biol 13, 1125–1135. 10.1016/j.chembiol.2006.09.009.

7. Ge, J., and Yu, Y.T. (2013). RNA pseudouridylation: new insights into an old modification. Trends Biochem Sci 38, 210–218. 10.1016/j.tibs.2013.01.002.

8. Borchardt, E.K., Martinez, N.M., and Gilbert, W.V. (2020). Regulation and Function of RNA Pseudouridylation in Human Cells. Annual Review of Genetics 54, 309–336. 10.1146/annurev-genet-112618-043830.

9. Lafontaine, D.L.J., and Tollervey, D. (1998). Birth of the snoRNPs: the evolution of the modification-guide snoRNAs. Trends in Biochemical Sciences 23, 383–388. 10.1016/S0968-0004(98)01260-2.

10. Garus, A., and Autexier, C. (2021). Dyskerin: an essential pseudouridine synthase with multifaceted roles in ribosome biogenesis, splicing, and telomere maintenance. Rna 27, 1441–1458. 10.1261/rna.078953.121.

11. Yoon, A., Peng, G., Brandenburg, Y., Zollo, O., Xu, W., Rego, E., and Ruggero, D. (2006). Impaired Control of IRES-Mediated Translation in X-Linked Dyskeratosis Congenita. Science 312, 902–906. doi:10.1126/science.1123835.

12. Bellodi, C., Kopmar, N., and Ruggero, D. (2010). Deregulation of oncogene-induced senescence and p53 translational control in X-linked dyskeratosis congenita. The EMBO Journal 29, 1865–1876. 10.1038/emboj.2010.83.

13. Montanaro, L., Calienni, M., Bertoni, S., Rocchi, L., Sansone, P., Storci, G., Santini, D., Ceccarelli, C., Taffurelli, M., Carnicelli, D., et al. (2010). Novel Dyskerin-Mediated Mechanism of p53 Inactivation through Defective mRNA Translation. Cancer Research 70, 4767–4777. 10.1158/0008-5472.Can-09-4024.

14. Jack, K., Bellodi, C., Landry, Dori M., Niederer, Rachel O., Meskauskas, A., Musalgaonkar, S., Kopmar, N., Krasnykh, O., Dean, Alison M., Thompson, Sunnie R., et al. (2011). rRNA Pseudouridylation Defects Affect Ribosomal Ligand Binding and Translational Fidelity from Yeast to Human Cells. Molecular Cell 44, 660–666. 10.1016/j.molcel.2011.09.017.

15. Montanaro, L. (2010). Dyskerin and cancer: more than telomerase. The defect in mRNA translation helps in explaining how a proliferative defect leads to cancer. The Journal of Pathology 222, 345–349. 10.1002/path.2777.

16. Alawi, F., and Lee, M.N. (2007). DKC1 is a direct and conserved transcriptional target of c-MYC. Biochemical and Biophysical Research Communications 362, 893–898. 10.1016/j.bbrc.2007.08.071.

17. O’Brien, R., Tran, S.L., Maritz, M.F., Liu, B., Kong, C.F., Purgato, S., Yang, C., Murray, J., Russell, A.J., Flemming, C.L., et al. (2016). MYC-Driven Neuroblastomas Are Addicted to a Telomerase-Independent Function of Dyskerin. Cancer Research 76, 3604–3617. 10.1158/0008-5472.Can-15-0879.

18. Meyer, N., and Penn, L.Z. (2008). Reflecting on 25 years with MYC. Nat Rev Cancer 8, 976–990. 10.1038/nrc2231.

19. Dang, C.V. (2012). MYC on the path to cancer. Cell 149, 22–35. 10.1016/j.cell.2012.03.003.

20. Rickman, D.S., Schulte, J.H., and Eilers, M. (2018). The Expanding World of N-MYC-Driven Tumors. Cancer Discov 8, 150–163. 10.1158/2159-8290.Cd-17-0273.

21. Bansal, M., Gupta, A., and Ding, H.-F. (2022). MYCN and Metabolic Reprogramming in Neuroblastoma. Cancers 14, 4113.

22. Cohn, S.L., Pearson, A.D.J., London, W.B., Monclair, T., Ambros, P.F., Brodeur, G.M., Faldum, A., Hero, B., Iehara, T., Machin, D., et al. (2009). The International Neuroblastoma Risk Group (INRG) Classification System: An INRG Task Force Report. J Clin Oncol 27, 289–297. 10.1200/jco.2008.16.6785.

23. Qiu, B., and Matthay, K.K. (2022). Advancing therapy for neuroblastoma. Nat Rev Clin Oncol 19, 515–533. 10.1038/s41571-022-00643-z.

24. Ryoo, H.D. (2024). The integrated stress response in metabolic adaptation. Journal of Biological Chemistry 300, 107151. 10.1016/j.jbc.2024.107151.

25. Kilberg, M.S., Balasubramanian, M., Fu, L., and Shan, J. (2012). The transcription factor network associated with the amino acid response in mammalian cells. Advances in nutrition 3, 295–306. 10.3945/an.112.001891.

26. Pakos-Zebrucka, K., Koryga, I., Mnich, K., Ljujic, M., Samali, A., and Gorman, A.M. (2016). The integrated stress response. EMBO reports 17, 1374–1395. 10.15252/embr.201642195.

27. Wortel, I.M.N., van der Meer, L.T., Kilberg, M.S., and van Leeuwen, F.N. (2017). Surviving Stress: Modulation of ATF4-Mediated Stress Responses in Normal and Malignant Cells. Trends Endocrinol Metab 28, 794–806. 10.1016/j.tem.2017.07.003.

28. Zhang, W., Yu, Y., Hertwig, F., Thierry-Mieg, J., Zhang, W., Thierry-Mieg, D., Wang, J., Furlanello, C., Devanarayan, V., Cheng, J., et al. (2015). Comparison of RNA-seq and microarray-based models for clinical endpoint prediction. Genome Biol 16, 133. 10.1186/s13059-015-0694-1.

29. Rajbhandari, P., Lopez, G., Capdevila, C., Salvatori, B., Yu, J., Rodriguez-Barrueco, R., Martinez, D., Yarmarkovich, M., Weichert-Leahey, N., Abraham, B.J., et al. (2018). Cross-Cohort Analysis Identifies a TEAD4-MYCN Positive Feedback Loop as the Core Regulatory Element of High-Risk Neuroblastoma. Cancer Discov 8, 582–599. 10.1158/2159-8290.Cd-16-0861.

30. Rocchi, L., Barbosa, A.J.M., Onofrillo, C., Del Rio, A., and Montanaro, L. (2014). Inhibition of Human Dyskerin as a New Approach to Target Ribosome Biogenesis. PLOS ONE 9, e101971. 10.1371/journal.pone.0101971.

31. Kan, G., Wang, Z., Sheng, C., Chen, G., Yao, C., Mao, Y., and Chen, S. (2021). Dual Inhibition of DKC1 and MEK1/2 Synergistically Restrains the Growth of Colorectal Cancer Cells. Advanced Science 8, 2004344. 10.1002/advs.202004344.

32. Thiaville, M.M., Dudenhausen, E.E., Zhong, C., Pan, Y.X., and Kilberg, M.S. (2008). Deprivation of protein or amino acid induces C/EBPbeta synthesis and binding to amino acid response elements, but its action is not an absolute requirement for enhanced transcription. Biochem J 410, 473–484. 10.1042/BJ20071252.

33. Hansen, B.S., Vaughan, M.H., and Wang, L. (1972). Reversible inhibition by histidinol of protein synthesis in human cells at the activation of histidine. J Biol Chem 247, 3854–3857.

34. Pang, Z., Lu, Y., Zhou, G., Hui, F., Xu, L., Viau, C., Spigelman, Aliya F., MacDonald, Patrick E., Wishart, David S., Li, S., and Xia, J. (2024). MetaboAnalyst 6.0: towards a unified platform for metabolomics data processing, analysis and interpretation. Nucleic Acids Research. 10.1093/nar/gkae253.

35. Nwosu, Z.C., Song, M.G., di Magliano, M.P., Lyssiotis, C.A., and Kim, S.E. (2023). Nutrient transporters: connecting cancer metabolism to therapeutic opportunities. Oncogene 42, 711–724. 10.1038/s41388-023-02593-x.

36. Hyde, R., Taylor, P.M., and Hundal, H.S. (2003). Amino acid transporters: roles in amino acid sensing and signalling in animal cells. Biochem J 373, 1–18.

37. Xia, Y., Ye, B., Ding, J., Yu, Y., Alptekin, A., Thangaraju, M., Prasad, P.D., Ding, Z.-C., Park, E.J., Choi, J.-H., et al. (2019). Metabolic Reprogramming by MYCN Confers Dependence on the Serine-Glycine-One-Carbon Biosynthetic Pathway. Cancer Res 79, 3837–3850. 10.1158/0008-5472.Can-18-3541.

38. King, Helen A., Cobbold, Laura C., and Willis, Anne E. (2010). The role of IRES trans-acting factors in regulating translation initiation. Biochemical Society Transactions 38, 1581–1586. 10.1042/bst0381581.

39. Bellodi, C., Krasnykh, O., Haynes, N., Theodoropoulou, M., Peng, G., Montanaro, L., and Ruggero, D. (2010). Loss of Function of the Tumor Suppressor DKC1 Perturbs p27 Translation Control and Contributes to Pituitary Tumorigenesis. Cancer Research 70, 6026–6035. 10.1158/0008-5472.Can-09-4730.

40. Rocchi, L., Pacilli, A., Sethi, R., Penzo, M., Schneider, R.J., Treré, D., Brigotti, M., and Montanaro, L. (2013). Dyskerin depletion increases VEGF mRNA internal ribosome entry site-mediated translation. Nucleic Acids Research 41, 8308–8318. 10.1093/nar/gkt587.

41. Damiano, F., Rochira, A., Tocci, R., Alemanno, S., Gnoni, A., and Siculella, L. (2012). hnRNP A1 mediates the activation of the IRES-dependent SREBP-1a mRNA translation in response to endoplasmic reticulum stress. Biochemical Journal 449, 543–553. 10.1042/bj20120906.

42. Roy, R., Durie, D., Li, H., Liu, B.-Q., Skehel, J.M., Mauri, F., Cuorvo, Lucia V., Barbareschi, M., Guo, L., Holcik, M., et al. (2014). hnRNPA1 couples nuclear export and translation of specific mRNAs downstream of FGF-2/S6K2 signalling. Nucleic Acids Research 42, 12483–12497. 10.1093/nar/gku953.

43. Dickson, L.M., and Brown, A.J.P. (1998). mRNA translation in yeast during entry into stationary phase. Molecular and General Genetics MGG 259, 282–293. 10.1007/s004380050814.

44. Roy, R., Huang, Y., Seckl, M.J., and Pardo, O.E. (2017). Emerging roles of hnRNPA1 in modulating malignant transformation. Wiley Interdiscip Rev RNA 8. 10.1002/wrna.1431.

45. Gebauer, F., Schwarzl, T., Valcárcel, J., and Hentze, M.W. (2021). RNA-binding proteins in human genetic disease. Nature Reviews Genetics 22, 185–198. 10.1038/s41576-020-00302-y.

46. Van Nostrand, E.L., Freese, P., Pratt, G.A., Wang, X., Wei, X., Xiao, R., Blue, S.M., Chen, J.-Y., Cody, N.A.L., Dominguez, D., et al. (2020). A large-scale binding and functional map of human RNA-binding proteins. Nature 583, 711–719. 10.1038/s41586-020-2077-3.

47. Jo, O.D., Martin, J., Bernath, A., Masri, J., Lichtenstein, A., and Gera, J. (2008). Heterogeneous Nuclear Ribonucleoprotein A1 Regulates Cyclin D1 and c-myc Internal Ribosome Entry Site Function through Akt Signaling*. Journal of Biological Chemistry 283, 23274–23287. 10.1074/jbc.M801185200.

48. Godet, A.-C., David, F., Hantelys, F., Tatin, F., Lacazette, E., Garmy-Susini, B., and Prats, A.-C. (2019). IRES Trans-Acting Factors, Key Actors of the Stress Response. International journal of molecular sciences 20, 924.

49. Bonnal, S., Pileur, F., Orsini, C., Parker, F., Pujol, F., Prats, A.-C., and Vagner, S. (2005). Heterogeneous Nuclear Ribonucleoprotein A1 Is a Novel Internal Ribosome Entry Site trans-Acting Factor That Modulates Alternative Initiation of Translation of the Fibroblast Growth Factor 2 mRNA*. Journal of Biological Chemistry 280, 4144–4153. 10.1074/jbc.M411492200.

50. Chan, C.-P., Kok, K.-H., Tang, H.-M.V., Wong, C.-M., and Jin, D.-Y. (2013). Internal ribosome entry site-mediated translational regulation of ATF4 splice variant in mammalian unfolded protein response. Biochimica et Biophysica Acta (BBA) - Molecular Cell Research 1833, 2165–2175. 10.1016/j.bbamcr.2013.05.002.

51. Taoka, M., Nobe, Y., Yamaki, Y., Sato, K., Ishikawa, H., Izumikawa, K., Yamauchi, Y., Hirota, K., Nakayama, H., Takahashi, N., and Isobe, T. (2018). Landscape of the complete RNA chemical modifications in the human 80S ribosome. Nucleic Acids Research 46, 9289–9298. 10.1093/nar/gky811.

52. Marchand, V., Pichot, F., Neybecker, P., Ayadi, L., Bourguignon-Igel, V., Wacheul, L., Lafontaine, D.L.J., Pinzano, A., Helm, M., and Motorin, Y. (2020). HydraPsiSeq: a method for systematic and quantitative mapping of pseudouridines in RNA. Nucleic Acids Research 48, e110–e110. 10.1093/nar/gkaa769.

53. Xu, H., Kong, L., Cheng, J., Al Moussawi, K., Chen, X., Iqbal, A., Wing, P.A.C., Harris, J.M., Tsukuda, S., Embarc-Buh, A., et al. (2024). Absolute quantitative and base-resolution sequencing reveals comprehensive landscape of pseudouridine across the human transcriptome. Nature Methods 21, 2024–2033. 10.1038/s41592-024-02439-8.

54. Zhang, M., Jiang, Z., Ma, Y., Liu, W., Zhuang, Y., Lu, B., Li, K., Peng, J., and Yi, C. (2023). Quantitative profiling of pseudouridylation landscape in the human transcriptome. Nature Chemical Biology. 10.1038/s41589-023-01304-7.

55. Bergeron, D., Paraqindes, H., Fafard-Couture, É., Deschamps-Francoeur, G., Faucher-Giguère, L., Bouchard-Bourelle, P., Abou Elela, S., Catez, F., Marcel, V., and Scott, Michelle S. (2022). snoDB 2.0: an enhanced interactive database, specializing in human snoRNAs. Nucleic Acids Research 51, D291–D296. 10.1093/nar/gkac835.

56. Tafer, H., Kehr, S., Hertel, J., Hofacker, I.L., and Stadler, P.F. (2009). RNAsnoop: efficient target prediction for H/ACA snoRNAs. Bioinformatics 26, 610–616. 10.1093/bioinformatics/btp680.

57. Zacchini, F., Venturi, G., De Sanctis, V., Bertorelli, R., Ceccarelli, C., Santini, D., Taffurelli, M., Penzo, M., Treré, D., Inga, A., et al. (2022). Human dyskerin binds to cytoplasmic H/ACA-box-containing transcripts affecting nuclear hormone receptor dependence. Genome Biology 23, 177. 10.1186/s13059-022-02746-3.

58. Faucher-Giguère, L., Roy, A., Deschamps-Francoeur, G., Couture, S., Nottingham, R.M., Lambowitz, A.M., Scott, M.S., and Abou Elela, S. (2022). High-grade ovarian cancer associated H/ACA snoRNAs promote cancer cell proliferation and survival. NAR Cancer 4. 10.1093/narcan/zcab050.

59. Kilberg, M.S., Shan, J., and Su, N. (2009). ATF4-dependent transcription mediates signaling of amino acid limitation. Trends Endocrinol Metab 20, 436–443. 10.1016/j.tem.2009.05.008.

60. Schmidt, S., Gay, D., Uthe, F.W., Denk, S., Paauwe, M., Matthes, N., Diefenbacher, M.E., Bryson, S., Warrander, F.C., Erhard, F., et al. (2019). A MYC–GCN2–eIF2α negative feedback loop limits protein synthesis to prevent MYC-dependent apoptosis in colorectal cancer. Nature Cell Biology 21, 1413–1424. 10.1038/s41556-019-0408-0.

61. Nguyen, H.G., Conn, C.S., Kye, Y., Xue, L., Forester, C.M., Cowan, J.E., Hsieh, A.C., Cunningham, J.T., Truillet, C., Tameire, F., et al. (2018). Development of a stress response therapy targeting aggressive prostate cancer. Science Translational Medicine 10, eaar2036. doi:10.1126/scitranslmed.aar2036.

62. Tameire, F., Verginadis, II, Leli, N.M., Polte, C., Conn, C.S., Ojha, R., Salas Salinas, C., Chinga, F., Monroy, A.M., Fu, W., et al. (2019). ATF4 couples MYC-dependent translational activity to bioenergetic demands during tumour progression. Nat Cell Biol 21, 889–899. 10.1038/s41556-019-0347-9.

63. Ding, J., Bansal, M., Cao, Y., Ye, B., Mao, R., Gupta, A., Sudarshan, S., and Ding, H.-F. (2024). MYC Drives mRNA Pseudouridylation to Mitigate Proliferation-Induced Cellular Stress during Cancer Development. Cancer Research 84, 4031–4048. 10.1158/0008-5472.Can-24-1102.

64. Genuth, N.R., and Barna, M. (2018). The Discovery of Ribosome Heterogeneity and Its Implications for Gene Regulation and Organismal Life. Molecular Cell 71, 364–374. 10.1016/j.molcel.2018.07.018.

65. Zhao, Y., Rai, J., and Li, H. (2023). Regulation of translation by ribosomal RNA pseudouridylation. Science Advances 9, eadg8190. doi:10.1126/sciadv.adg8190.

66. Harvey, R.F., Pöyry, T., and Willis, A.E. (2024). Let’s (P-s)talk about specialized ribosomes. Molecular Cell 84, 4478–4479. 10.1016/j.molcel.2024.11.011.

67. Tidu, A., and Martin, F. (2024). The interplay between cis- and trans-acting factors drives selective mRNA translation initiation in eukaryotes. Biochimie 217, 20–30. 10.1016/j.biochi.2023.09.017.

68. Spriggs, K.A., Stoneley, M., Bushell, M., and Willis, A.E. (2008). Re-programming of translation following cell stress allows IRES-mediated translation to predominate. Biology of the Cell 100, 27–38. 10.1042/BC20070098.

69. Komar, A.A., and Hatzoglou, M. (2011). Cellular IRES-mediated translation. Cell cycle (Georgetown, Tex 10, 229–240. 10.4161/cc.10.2.14472.

70. Harding, H.P., Zhang, Y., Zeng, H., Novoa, I., Lu, P.D., Calfon, M., Sadri, N., Yun, C., Popko, B., Paules, R., et al. (2003). An integrated stress response regulates amino acid metabolism and resistance to oxidative stress. Mol Cell 11, 619–633.

71. Costa-Mattioli, M., and Walter, P. (2020). The integrated stress response: From mechanism to disease. Science 368, eaat5314. doi:10.1126/science.aat5314.

72. Kovalski, J.R., Kuzuoglu-Ozturk, D., and Ruggero, D. (2022). Protein synthesis control in cancer: selectivity and therapeutic targeting. The EMBO Journal 41, e109823. 10.15252/embj.2021109823.

73. Spriggs, K.A., Bushell, M., and Willis, A.E. (2010). Translational Regulation of Gene Expression during Conditions of Cell Stress. Molecular Cell 40, 228–237. 10.1016/j.molcel.2010.09.028.

74. Marques, R., Lacerda, R., and Romão, L. (2022). Internal Ribosome Entry Site (IRES)-Mediated Translation and Its Potential for Novel mRNA-Based Therapy Development. Biomedicines 10, 1865.

75. Biedler, J.L., Roffler-Tarlov, S., Schachner, M., and Freedman, L.S. (1978). Multiple neurotransmitter synthesis by human neuroblastoma cell lines and clones. Cancer Res 38, 3751–3757.

76. Levengood, J.D., Potoyan, D., Penumutchu, S., Kumar, A., Zhou, Q., Wang, Y., Hansen, A.L., Kutluay, S., Roche, J., and Tolbert, B.S. (2024). Thermodynamic coupling of the tandem RRM domains of hnRNP A1 underlie its pleiotropic RNA binding functions. Science Advances 10, eadk6580. doi:10.1126/sciadv.adk6580.

77. Park, Y., Reyna-Neyra, A., Philippe, L., and Thoreen, C.C. (2017). mTORC1 Balances Cellular Amino Acid Supply with Demand for Protein Synthesis through Post-transcriptional Control of ATF4. Cell Rep 19, 1083–1090. 10.1016/j.celrep.2017.04.042.

78. Alptekin, A., Ye, B., Yu, Y., Poole, C.J., van Riggelen, J., Zha, Y., and Ding, H.-F. (2019). Glycine decarboxylase is a transcriptional target of MYCN required for neuroblastoma cell proliferation and tumorigenicity. Oncogene 38, 7504–7520. 10.1038/s41388-019-0967-3.

79. Ushmorov, A., Hogarty, M.D., Liu, X., Knauss, H., Debatin, K.M., and Beltinger, C. (2008). N-myc augments death and attenuates protective effects of Bcl-2 in trophically stressed neuroblastoma cells. Oncogene 27, 3424–3434. 10.1038/sj.onc.1211017.

80. Koster, J., Volckmann, R., Zwijnenburg, D., Molenaar, P., and Versteeg, R. (2019). Abstract 2490: R2: Genomics analysis and visualization platform. Cancer Research 79, 2490–2490. 10.1158/1538-7445.Am2019-2490.

81. Beg, A.A., Finco, T.S., Nantermet, P.V., and Baldwin, A.S., Jr. (1993). Tumor necrosis factor and interleukin-1 lead to phosphorylation and loss of I kappa B alpha: a mechanism for NF-kappa B activation. Mol Cell Biol 13, 3301–3310.

82. Lal, A., Mazan-Mamczarz, K., Kawai, T., Yang, X., Martindale, J.L., and Gorospe, M. (2004). Concurrent versus individual binding of HuR and AUF1 to common labile target mRNAs. The EMBO Journal 23, 3092–3102. 10.1038/sj.emboj.7600305.

83. Bremer, J., and van den Akker, P.C. (2022). In Vivo Models for the Evaluation of Antisense Oligonucleotides in Skin. Methods Mol Biol 2434, 315–320. 10.1007/978-1-0716-2010-6_21.

84. Bansal, M., Kundu, A., Gibson, A., Gupta, A., Ding, J., RudraRaju, S.V., Sudarshan, S., and Ding, H.-F. (2024). Transcriptome-wide quantitative profiling of PUS7-dependent pseudouridylation by nanopore direct long read RNA sequencing. bioRxiv, 2024.2001.2031.578250. 10.1101/2024.01.31.578250.

85. Li, H. (2018). Minimap2: pairwise alignment for nucleotide sequences. Bioinformatics 34, 3094–3100. 10.1093/bioinformatics/bty191.

86. Chen, Y., Sim, A., Wan, Y.K., Yeo, K., Lee, J.J.X., Ling, M.H., Love, M.I., and Göke, J. (2023). Context-aware transcript quantification from long-read RNA-seq data with Bambu. Nat Methods 20, 1187–1195. 10.1038/s41592-023-01908-w.

87. Hertz, M.I., Landry, D.M., Willis, A.E., Luo, G., and Thompson, S.R. (2013). Ribosomal Protein S25 Dependency Reveals a Common Mechanism for Diverse Internal Ribosome Entry Sites and Ribosome Shunting. Molecular and Cellular Biology 33, 1016–1026. 10.1128/MCB.00879-12.

88. Chen, H., and Boutros, P.C. (2011). VennDiagram: a package for the generation of highly-customizable Venn and Euler diagrams in R. BMC bioinformatics 12, 35. 10.1186/1471-2105-12-35.

